# α-Synuclein aggregation intermediates form fibril polymorphs with distinct prion-like properties

**DOI:** 10.1101/2020.05.03.074765

**Authors:** Surabhi Mehra, Sahil Ahlawat, Harish Kumar, Nitu Singh, Ambuja Navalkar, Komal Patel, Pradeep Kadu, Rakesh Kumar, Narendra N. Jha, Jayant B. Udgaonkar, Vipin Agarwal, Samir K. Maji

## Abstract

α-Synuclein (α-Syn) amyloid fibrils in synucleinopathies (such as Parkinson’s disease (PD), multiple system atrophy (MSA)) are structurally and functionally different, reminiscent of prion-like strains. However, how a single protein can form different fibril polymorphs in various synucleinopathies is not known. Here, we demonstrate the structure-function relationship of two distinct α-Syn fibril polymorphs, the pre-matured fibrils (PMF) and helix-matured fibrils (HMF) based on α-Syn aggregation intermediates. These polymorphs not only display the structural differences, including their fibril core structure as demonstrated by solid-state nuclear magnetic resonance (NMR) spectroscopy and H/D-exchange coupled with mass spectrometry but also possess different cellular activities such as seeding, cellular internalization, and cell-to-cell transmission. The HMF with a compact core structure exhibits low seeding potency *in cells* but readily internalizes and transmits from one cell to another. Whereas the less structured PMF lacks the cell-to-cell transmission ability but induces abundant α-Syn pathology and triggers the formation of aggresomes *in cells*. Overall, the study highlights how the conformational heterogeneity in the aggregation pathway may lead to fibril polymorphs with distinct prion-like behavior in PD.

## Introduction

Synucleinopathies are the group of neurological disorders, which are characterized by the presence of intracellular inclusion bodies composed of α-synuclein (α-Syn) amyloid fibrils^1^. Although α-Syn amyloids in neuronal inclusions termed as Lewy bodies (LBs) and Lewy neurites (LNs) are the characteristic feature of Parkinson’s disease (PD), the α-Syn inclusions in oligodendrocytes (glial cell inclusions; GCIs) are predominant in multiple system atrophy (MSA). Previous studies have suggested that amyloid fibrils associated with various neurodegenerative disorders are infectious and exhibit ‘prion-like’ behavior^2,3^. For example, exogenously added α-Syn fibrils readily internalize *in cells* and induce the aggregation of endogenous soluble α-Syn into pathogenic insoluble LB-like inclusions^4,5^. Moreover, the LB/LN-like inclusions are also observed in animals upon receiving intracerebral injections of synthetic α-Syn fibrils or brain homogenates from old transgenic (Tg) mice with α-Syn pathology^6,7^. This suggests prion-like cellular transmission and propagation of α-Syn amyloids in PD and related disorders^8^. In this context, it has been hypothesized that α-Syn assembles into polymorphs, which may account for distinct disease phenotypes observed amongst the synucleinopathies. α-Syn fibrils from GCIs in oligodendrocytes and LBs in neurons of diseased brain samples differ significantly in their structure and exhibit distinct seeding propensity^9^. Although α-Syn fibril polymorphs mostly differ in their fibril diameter, presence of twists, and the number of protofilaments^10,11^, they share a common cross-β-sheet structure with different packing and inter-protofilament interface^12,13^ as shown by cryo-EM. Previously, various experimental conditions have been used to develop α-Syn fibrils polymorphs *in vitro* towards establishing their structure-pathogenic relationship^14–17^. These polymorphs often result in distinct clinical and pathological phenotypes in the animal models^18,19^.

Since α-Syn amyloid formation involves a heterogeneous population of intermediate species^20–22^, we hypothesized that these intermediates species are capable of forming α-Syn polymorphs with distinct prion-like properties. Based on α-Syn aggregation intermediates, we developed two fibrillar polymorphs, PMF and HMF, with distinct fibril packing, structure and prion-like properties. HMF is more compact with a stable fibril core, readily internalizes in different cells and shows a higher tendency for trans-cellular spread. In contrast, PMF, without a robust protease-resistant fibril core, lacks the trans-cellular spreading ability but shows high seeding potency both *in vitro* and *in cells*. The present data support that the relative population of intermediate species might dictate the property of fibrils polymorphs and their biological activities associated with pathological traits in synucleinopathies.

## Results

### Isolation of different intermediates during aggregation and the subsequent generation of α-Syn polymorphs

α-Syn aggregation involves a helix-rich intermediate state at the nucleation step before converting to β-sheet-rich amyloid^23^. This helix-rich state constitutes a heterogeneous mixture of helical oligomers, β-sheet rich pre-matured fibrils, and soluble monomeric α-Syn. We proposed to develop α-Syn fibril polymorphs based on the intermediate species formed in this helix-rich state. For this, we incubated α-Syn low molecular weight (LMW) protein and monitored the aggregation kinetics by amyloid-specific Thioflavin T (ThT) dye (Figure S1a) and circular dichroism (CD) at regular intervals (Figure S1b). As soon as the helical conformation was observed in CD, we isolated different components based on the previously published protocol^23^. The pre-matured fibrils obtained as the by-product during isolation was termed as ‘PMF’ while the isolated helix was incubated further to form helix-matured fibrils (HMF) (Figure 1a, S1c). In a control set, LMW α-Syn was incubated and allowed to form fibrils, termed as Syn fibrils (Figure 1a, S1c). Throughout the study, the properties of PMF and HMF were compared to Syn fibrils (used as a control).

**Figure 1.**
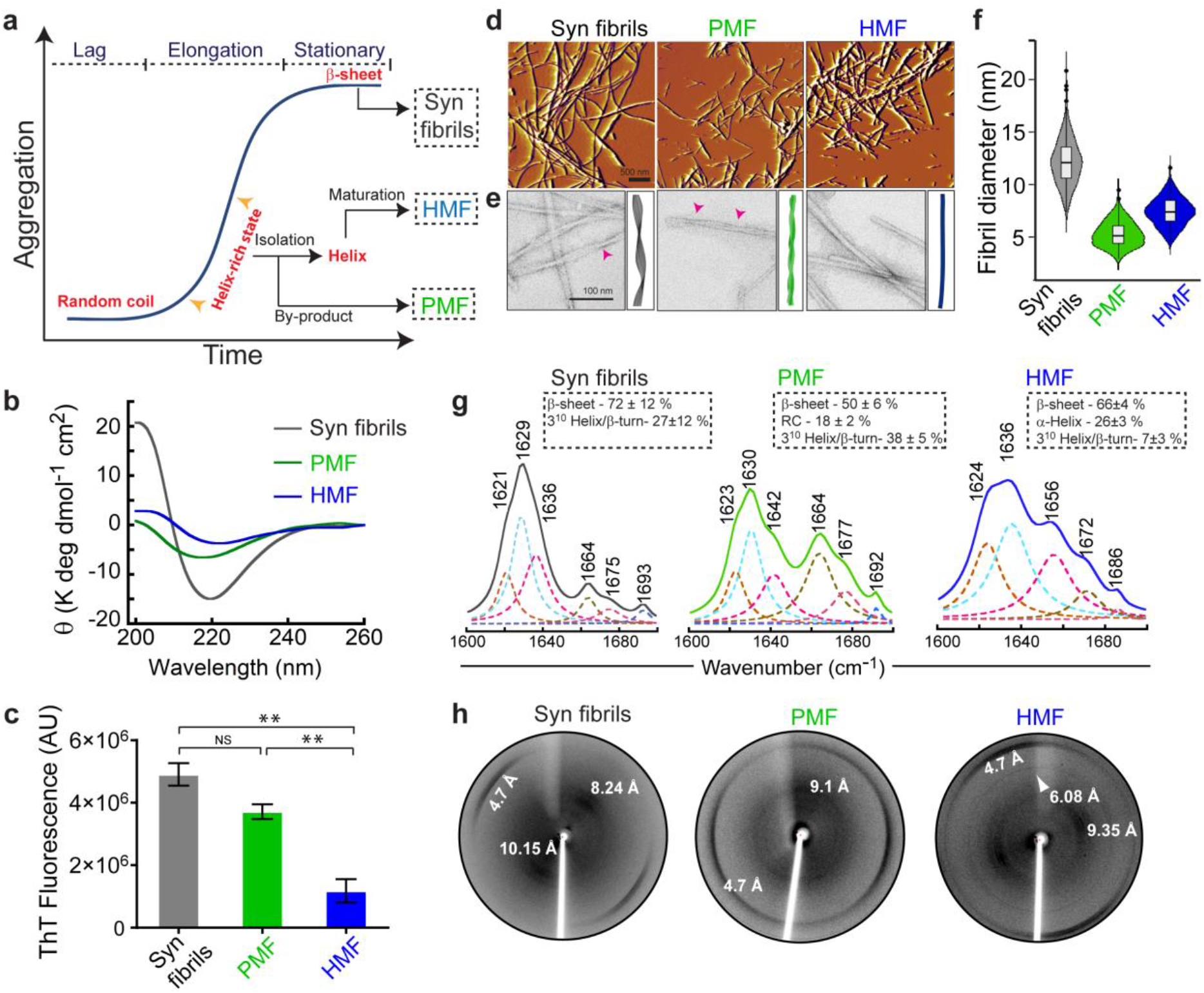
Isolation and biophysical characterization of α-Syn fibrils polymorphs. (a) Schematic for the isolation of intermediate species formed during α-Syn aggregation and subsequent generation of α-Syn polymorphs. (b) CD spectra of fibril polymorphs along with Syn fibrils, recorded at identical protein concentration shows a β-sheet structure with a single minimum at ~218-220 nm. (c) Thioflavin T fluorescence intensity after binding to different fibrils at 480 nm shows the highest fluorescence for Syn fibrils, followed by PMF and HMF. Morphology of the polymorphs and Syn fibrils visualized by (d) AFM and (e) TEM. The TEM images are shown along with the pictorial representations of the fibrils highlighting the morphological differences amongst different fibrils. Pink arrowheads mark the presence of twists and turns in Syn fibrils and PMF. Scale bar is 100 nm and 500 nm in TEM and AFM images, respectively. (f) Violin plot representing the distribution of fibril diameter analyzed from TEM images by using ImageJ software (n>250 counts). (g) Curve-fitted FTIR spectra in the region corresponding to the amide-I band of α-Syn fibrils (grey), PMF (green) and HMF (blue) along with Fourier deconvolution (dotted lines) are presented. The insets showing the percentage of secondary structural components in each case calculated from FTIR deconvolution. (h) X-ray diffraction pattern of the fibril variants. Fibrils diffraction pattern of Syn fibrils and PMF is typical of amyloids with a meridional peak at ~4.7Å with varied equatorial reflection. HMF shows multiple equatorial reflections along with an additional peak at 6.08 Å. Values in (c) and (g) represent mean ± s.e.m and mean ± s.d, respectively, n=3 from independent experiments. The statistical significance determined by one-way ANOVA followed by Newman-Keuls Multiple Comparison post hoc test; **P<0.01; NS P>0.05.

Post maturation, all three fibrillar variants showed a single minimum at ~218 nm in CD (Figure 1b) and bound to ThT dye suggesting the presence of a β-sheet rich structure (Figure 1c). The difference in CD signals at 218 nm, along with the different extent of ThT binding, suggests that these fibrils might possess different fibril packing and/or β-sheet conformation. Morphological analysis of fibrils by atomic force microscopy (AFM) and transmission electron microscopy (TEM) further revealed differences in the fibril diameter, presence of sub-filaments, and twisting behavior (Figure 1d, e and f). PMF formed thin fibrils with an average diameter of ~5 nm and consisted of 2-3 sub-filaments twisted around each other and also displayed high lateral association. In contrast, HMF was mostly untwisted with an average diameter of ~7 nm. However, both PMF and HMF were considerably thinner and shorter compared to Syn fibrils, which were long un-branched fibrils with an average diameter of ~12 nm (Figure 1e and f). This observation was also verified with AFM imaging of these fibril polymorphs (Figure 1d).

### PMF and HMF exhibit distinct structural properties and fibril core conformation

We next assessed the structural properties of these fibril polymorphs by Fourier-transform infrared (FTIR) spectroscopy, X-ray diffraction (XRD) and proteinase K digestion assay. The FTIR data suggest 72 %, 50 % and 66 % β-sheet content (~1620-1640 cm^−1^) for Syn fibrils, PMF and HMF, respectively (Figure 1g). In addition to β-sheet, PMF showed 18 % random coil (1642 cm^−1^) and 38 % 3^10^ helix/β-turn (~1664-1677 cm^−1^) content; whereas HMF showed 26 % of helical content (1656 cm^−1^) (Figure 1g). FTIR analysis, therefore, suggests that both HMF and PMF have substantial differences in their secondary structural components. The structural differences between PMF and HMF were further apparent from XRD analysis. Although the Syn fibrils showed typical reflections for cross-β-sheet structure^24^ (meridional and equatorial arcs at 4.7 Å and ~8-10 Å, respectively), PMF showed diffused equatorial arc at 9 Å along with meridional arc at 4.6 Å (Figure 1h). The diffraction pattern of HMF, however, revealed a distinct fibrillar architect with an additional arc at ~6 Å along with typical cross β-sheet reflections (Figure 1h).

Amyloids formed by various proteins/peptides possess protease-resistant core^25,26^, similar to infectious scrapie prions^27^. Various biophysical and structural studies have shown that residues 31–109 of α-Syn form the fibril core^25,28^. This core region is buried within the amyloid backbone and is likely involved in the fibril assembly. To examine any possible differences in the protease-resistant core of the polymorphs, we employed time-dependent proteinase K (PK) digestion and analyzed the digested samples with SDS-PAGE and mass spectrometry. At early timepoints, all the fibrils showed a prominent PK-resistant band migrating at ~11 kDa along with multiple low-molecular-weight bands in SDS-PAGE (Figure 2a). Interestingly, post 15 mins of treatment, although Syn fibrils and HMF retained the protease-resistant core, it disappeared in PMF. This was confirmed with MALDI-TOF mass spectrometry analysis, which revealed major peaks close to ~ 10923, ~8845, and ~7265 Da (Figure 2b) for all three fibril variants till 30 mins of protease digestion, but subsequently disappeared in PMF over prolonged digestion (Figure 2b). These results suggest that the core region of PMF is possibly more solvent-exposed compared to the HMF and Syn fibrils.

**Figure 2.**
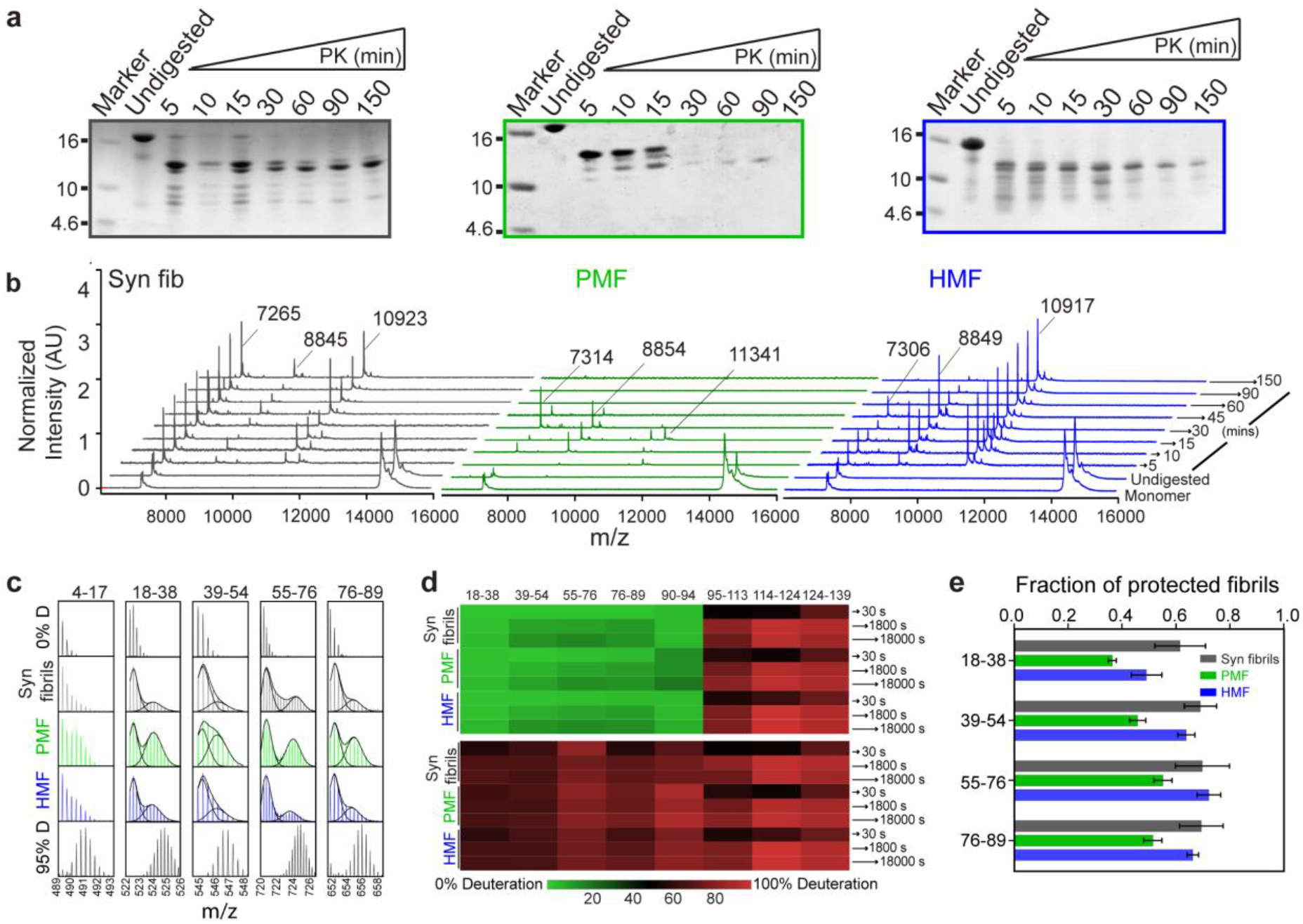
Characteristics difference in fibril core of HMF and PMF. (a-b) Proteinase K digestion profile of α-Syn polymorphs (100 μM) along with Syn fibrils monitored over time and analyzed by (a) Coomassie-stained SDS-PAGE (15%). Molecular weight ladder (kDa) is shown on the left of each gel. (b) MALDI-TOF mass spectrometry of the fibrils treated with 10 μg/ml Proteinase-K at each time point. (c-e) HDX-MS of α-Syn fibrillar polymorphs. (c) Representative mass spectra of peptide fragments of fibrillar variants along with protonated (0% D) and deuterated (95% D) spectra as controls. HDX-MS, performed at 1800 sec labeling pulse in deuterated buffer at pH 8.0, 37 °C is shown. Peptide fragments from 18-38, 39-54 and 55-76 shows a bimodal mass distribution for all fibril samples. (d) Heat map of % deuterium incorporation into different sequence-segments of fibrils. The sequence segments from 18-38, 39-54, 55-76, 76-89, and 90-94 shows two fibril populations; *upper panel*, one with lower mass (protected type P1) in observed binomial distribution and *lower panel*, showing population with higher mass (protected type P2) of a binomial distribution. The peptide fragments in the C-terminus region (95-139) gets fully labeled and show 80-100 % deuterium incorporation. (d) Bar plot represents the fraction of protected fibril (P1) in each polymorph (for residues 18-38, 39-54, 55-76 and 76-89) at 1800 seconds labeling pulse. Values represent mean ± s.e.m, n=3 from independent experiments.

Further to gain residue/segment-specific information for the possible differences in the fibril core of HMF and PMF, we performed HDX-MS analysis^29–31^. For all the fibril polymorphs, residues 4 to 95 remained highly protected (unlabeled similar to 0 % deuterated α-Syn monomer peaks) in contrast to C-terminal residues 96 to 140 (labeled similar to the 95 % deuterated monomer peaks) (Figure 2c, Figure S2 and S3). This suggests that residues 4 to 95 contain the inaccessible backbone amide hydrogens, while C-terminus (residues 96 to 140) is mostly disordered. Interestingly, the segments spanning residues 18 to 95 showed a bimodal mass distribution for all the fibrils (Figure 2c, Figure S3), indicating the existence of two different fibrillar conformations as observed previously for α-Syn oligomers and fibrils^21,31,32^. To quantify the relative population of the two conformations, the data were fitted to the sum of two Gaussian distributions. One of the conformations (P1) with a lower mass, in the observed binomial distribution, displayed protection against HDX, suggesting to be ordered and protected type (Figure 2d, *upper panel*). The other conformation (P2), with a higher mass, displayed no protection and is possibly unstructured (Figure 2d, *lower panel*). The amount of the solvent protected population in the peptides with residues 18-38, 39-54, 76-89, and 90-94 differed significantly for HMF and PMF polymorphs (Figure 2e). Similar to Syn fibrils (residues 4-95), the residues 39-89 of HMF showed ~67 % protected population, indicating that HMF is mostly ordered and structured fibril type. On the contrary, PMF (residues 4-95) displayed only ~40 % solvent protected population (Figure 2e). The HDX-MS results are in agreement with PK digestion assay, thus indicating PMF to be a flexible and relatively unordered fibril type compared to HMF.

### Structural differences between α-Syn fibril polymorphs revealed by solid-state NMR spectroscopy (ssNMR)

We employed ssNMR to characterize the local changes in structures at atomic resolution amongst fibril polymorphs. To probe residue-level differences between fibrils, backbone atoms were assigned using two-dimensional (DARR, NCA) and three-dimensional (NCACB, NCACX, CANCO, NCOCX) with U-[^13^C,^15^N] labeled samples of fibril polymorphs^33^. For HMF and Syn fibrils, the first 80 and 72 residues out of the starting 98 residues were assigned, respectively (Figure S4). Similar to previous reports^34^, we observed that for these two fibril forms, the region spanning residues 98-140 was flexible and could not be observed in the cross-polarization (CP) based experiments (Figure 3, S4 and S5). For HMF, the flexibility of these residues was independently confirmed by INEPT experiments^35^. We further observed that the spectra for HMF and Syn fibrils were similar (Figure 3a and S4) however not identical, as indicated by the plot of the chemical shift differences in the backbone atom (Figure S4a). The differences in chemical shift values of residues were majorly present in the region from 40-60 (Figure 3a, S4). Due to difficulty in preparation of labeled PMF in larger quantities, required to perform ^13^C detected multidimensional experiment, a residue-specific assignment on the PMF sample was not performed. However, the structural comparison between PMF and HMF/Syn fibrils was drawn based on 2D ^13^C-^13^C and ^15^N-^13^C spectra recorded under identical conditions (Figure 3b and c, S5). Due to the absence of atomic level assignment for PMF, only the well-resolved glycine, threonine, and serine regions of the spectra were compared for the three fibrils forms. The independent overlay of PMF with the HMF (Figure 3b and S5a) and Syn fibril spectra (Figure 3c and S5b) suggest that most peak positions were conserved, barring minor differences within the experimental limits. The major differences between the PMF and the other two-fibril forms appear to be in the residues within the non-amyloid-β component (NAC) region (61-95). In the PMF sample, the peaks corresponding to 73G and 75T were completely absent (Figure 3b and c, zoom-in spectra; red and blue circles) in contrast to the other two fibril forms. The cross peak due to 72T was very weak (Figure S5), and the other residues in the segment 72-78 are alanine and valine, which were difficult to resolve solely based on 2D spectra. When compared with HMF, these cumulative changes suggest that a well-structured β-sheet in HMF from residues 74-79 segment is either absent or resides in an entirely different chemical or dynamic environment in PMF. Similarly, the overlay plot for the Syn fibrils and PMF samples revealed several minor differences in the chemical shifts. Significant changes appeared in the peak position for the nitrogen spin of 42S and peak positions of 41/47G and 44/54T (Figure 3c, blue dotted circles), thereby implying that the residues 40-60 are more ordered in the PMF sample compared to Syn fibrils. Similar to HMF polymorph, the most notable change between PMF and Syn fibrils is again the absence of 73G and 75T peaks (Figure 3c, zoom-in spectra), thereby implying differences in the NAC region.

**Figure 3.**
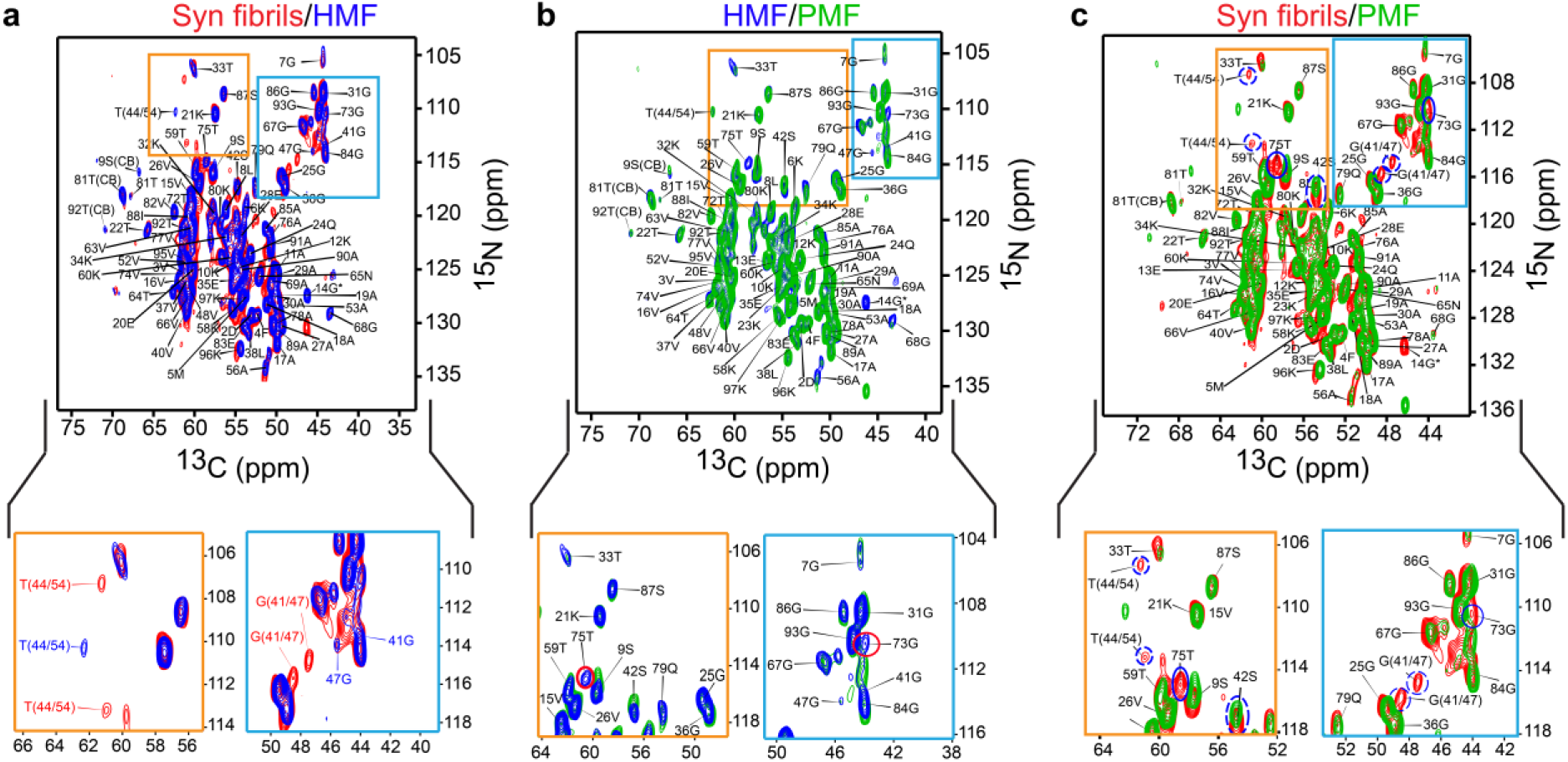
Solid-state NMR of α-Syn polymorphs. (a) *Upper panel* shows the overlay of 2D NCA spectra for uniformly ^13^C,^15^N labeled Syn fibrils (red) and HMF (blue). *Lower panel* shows the zoomed-in region marked by orange and blue boxes in the 2D NCA spectrum depicting residue-level differences. The assignments of Syn fibrils are marked with red and HMF in blue. (b) *Upper panel* shows the overlay of the 2D NCA spectra for uniformly ^13^C,^15^N labeled PMF (green) and HMF (blue) samples and *lower panel* shows the zoomed-in region marked by orange and blue boxes in the 2D NCA spectrum highlighting the differences in glycine and threonine regions between two-fibril polymorphs. Red circles highlight differences mentioned in text. (c) *Upper panel* shows the overlay of 2D NCA spectra of PMF (green) and Syn fibrils (red) samples. Lower panel represents the zoomed-in region marked by orange and blue boxes in 2D NCA spectrum. Solid blue circles mark the absence of 75T and 73G in PMF sample and dotted blue circles highlight differences in 42S and peak positions of 41/47G and 44/54T between two-fibril polymorphs. In (a) and (b) the assignments of only HMF while in (c) the assignments of only Syn fibrils are shown for clarity unless otherwise mentioned. The 14G peak (marked with *) in NCA spectra is folded back in all the spectra.

The differences between the fibril polymorphs thus appear to be in the two regions of the protein chain; (i) between residues 40-60 and (ii) in the NAC region. The HMF and Syn fibrils have a structurally identical N-terminal (1-40) and the C-terminus (61-140). The difference between the HMF and Syn fibril appears to be mainly in the 40-60 residues segment and arises due to structural heterogeneity. Notably, the NAC region in HMF and Syn fibrils is also identical, indicating a similar structure of the core. Intriguingly, the major difference in the PMF and the other two polymorphs is predominantly found in the NAC domain (60-80 residues) This structural difference in the NAC region could probably be the reason for the lack of proteinase K resistance and amide protection in case of PMF. From this investigation, it is unequivocally evident that the NAC region plays a defining role in actuating the biophysical and biochemical properties of the different polymorphs.

### PMF exhibits high seeding ability *in vitro*

Seeded protein aggregation is now a well-known mechanism for the amyloid fibril formation and underlies the basis of prion-like propagation of amyloids^36^. The seed eliminates the requirement of primary nucleation by acting as a template to accelerate the aggregation kinetics by two plausible mechanisms; elongation and surface-catalysis^37^. To test if the PMF/HMF can seed and also template the conversion of soluble species into fibrils (Figure 4a), we performed the aggregation kinetics of α-Syn in the presence of trace amounts of fibrils seeds. For quantifying the relative efficiency of seeding, we first optimized the sonication time and used pre-formed fibril seeds of equal length (~65 nm) for seeding experiments (Figure 4b and Figure S6a and b).

**Figure 4.**
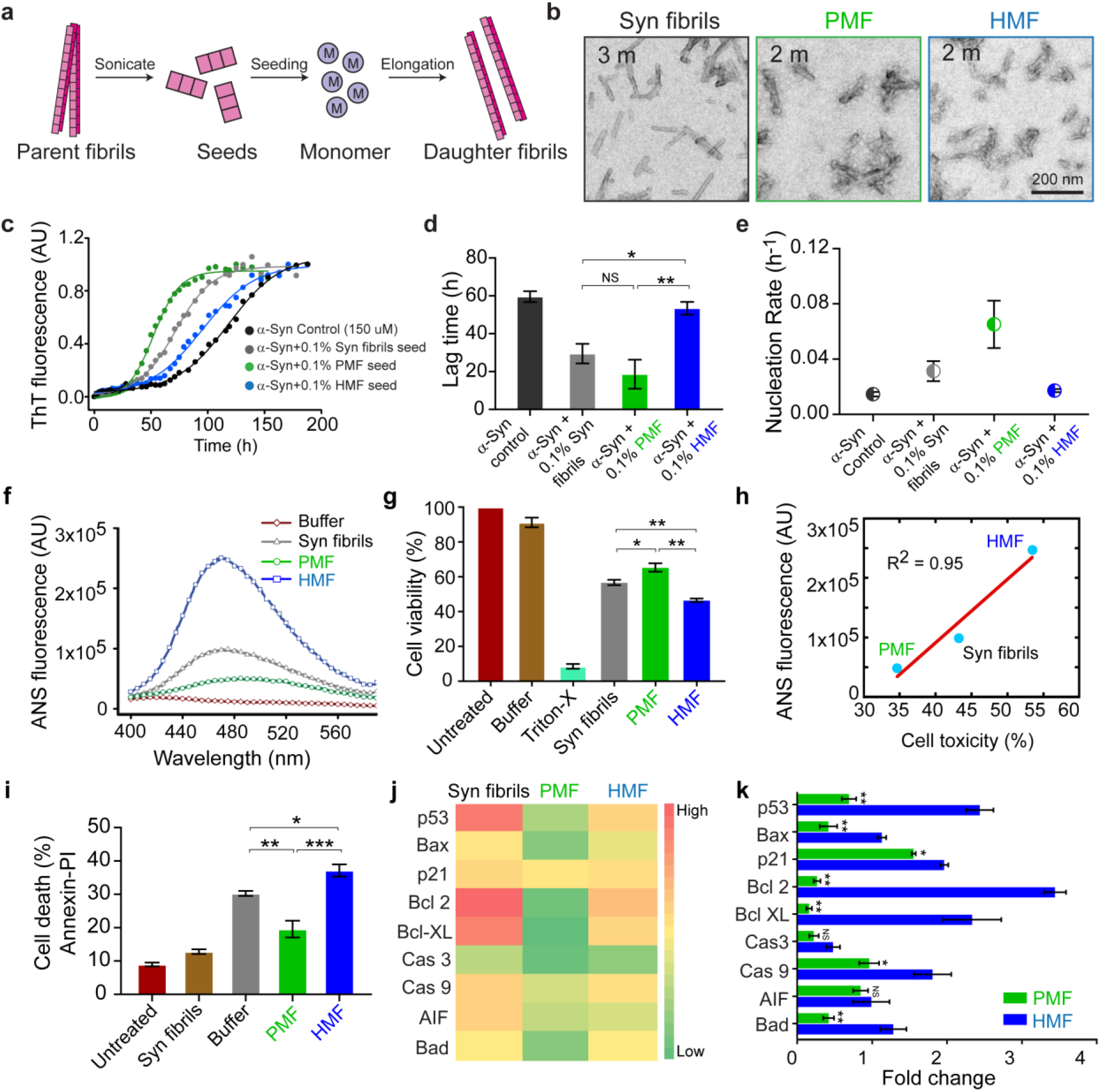
Seeding and toxicity of fibrillar polymorphs. (a) Schematic representation of the seeding and templating by pre-formed fibril seeds and generation of daughter fibrils. (b) TEM images of Syn fibrils, PMF and HMF after sonication at 3 mins, 2 mins, and 2 mins, respectively. These time points are optimized to obtain a similar length of fibril seeds. (c) Aggregation kinetics of α-Syn (150 μM protein concentration) in the presence of 0.1% pre-formed fibrillar seeds of PMF, HMF and Syn fibrils monitored by ThT assay in a microplate reader. α-Syn (150 μM) in the absence of fibril seeds was used as control. (d) The bar plot showing the lag time of aggregation and (e) nucleation rate obtained from the second derivate plots of the aggregation kinetics. (f) Hydrophobic surface exposure determined by ANS fluorescence of α-Syn polymorphs in the wavelength range of 400-600 nm. (g) The viability of SH-SY5Y cells treated with fibril polymorphs along with control samples for 24 h determined by MTT assay. HMF shows maximum cytotoxicity, followed by Syn fibrils and PMF. (h) Correlation plot of ANS fluorescence and cellular toxicity (R^2^= 0.95) using MTT assay showing that species with higher hydrophobic exposed surfaces are more toxic. (i) Cell-death analysis of SH-SY5Y cells treated with different fibrils by Annexin V-PI binding assay using flow cytometry. (j) Heat map of apoptotic gene profiling of SH-SY5Y cells treated with α-Syn fibrils, HMF and PMF under similar conditions used for toxicity assay by Annexin V-PI binding assay. The data is normalized with GAPDH and buffer was used as a control. Red color indicates higher and green color indicates lower gene expression with respect to buffer control as shown in the corresponding color map. (k) Bar plot of fold change expression in cells treated with HMF and PMF. The data demonstrate significant upregulation of apoptotic genes in cells treated with HMF compared to PMF. The values in (d), (e), (h), (i), and (k) represents mean ± s.e.m, n= 3 from independent experiments.*P≤0.05; **P≤0.01;***P≤0.001 NS P>0.05; determined by one-way ANOVA followed by Newman Keuls Multiple Comparison post hoc test in all the plots.

To test the seeding ability of the fibril polymorphs, we performed α-Syn aggregation (150 μM α-Syn monomer) in the presence of 0.1% (v/v) fibrillar seeds and monitored the kinetics by ThT binding assay. The aggregation of α-Syn showed a sigmoidal growth curve (Figure 4c). Upon addition of pre-formed fibrillar seeds, we observed acceleration of aggregation kinetics, as seen by a reduction in the lag phase (Figure 4c). We calculated the lag time by fitting the growth curve with the established equations^38^. α-Syn alone aggregated (in the absence of any seed) with a lag time ~ 60 h. PMF, however, showed a significant reduction in the lag time (~19 h), followed by Syn fibrils (~29 h). HMF, on the contrary, showed a lag time of ~53 h, which was significantly high compared to the other two fibril types (Figure 4d). The reduction in the lag time also correlated with the nucleation rate of the different polymorphs, where PMF showed the highest nucleation rate (0.065 h^−1^) compared to Syn fibrils (0.031 h^−1^) and HMF (0.017 h^−1^) (Figure 4e). Next, we analyzed the morphology of the aggregates formed after seeding and compared it with that of the parent fibrils. TEM images revealed that the morphological differences were retained even in the daughter fibrils generated from the seeded aggregation (Figure S6c). The data, therefore, suggest that although both PMF and HMF can seed the soluble monomeric species *in vitro*, the rate of recruitment of monomeric species and seeding potency is significantly different for the two polymorphs.

### Exposure of hydrophobic surfaces of fibril polymorphs results in different cytotoxicity

Since exposed hydrophobic surfaces of protein aggregates may dictate its cellular toxicity^39,40^, we analyzed hydrophobic surface exposure of all the fibril polymorphs using 1-anilinonaphthalene-8-sulfonate (ANS) and Nile red (NR) binding assay. HMF showed maximum ANS binding with nearly three and five-fold increase in fluorescence intensity compared to Syn fibrils and PMF, respectively (Figure 4f). A similar observation was also obtained from NR dye-binding assay (Figure S7a), suggesting that hydrophobic surfaces are considerably exposed in the case of HMF. Interestingly, the higher exposed hydrophobic surface of HMF makes it more cytotoxic compared to other fibril polymorphs as evident from MTT (4,5-Dimethylthiazol-2-yl)-2,5-diphenyltetrazoliumbromide) toxicity assay using human neuroblastoma SH-SY5Y cell line (Figure 4g and h). Cells treated with HMF and PMF showed 46.5 % and 65 % cell viability, respectively whereas Syn fibrils showed 57 % cell viability. To further quantify the toxicity measurement, cell death analysis was done by Annexin V and propidium iodide (PI) staining assay followed by flow cytometry analysis^41^. Consistent with MTT data (Figure 4g), HMF displayed maximum cytotoxicity (37 %) followed by Syn fibrils (30 %) in Annexin V-PI assay. However, cells treated with PMF were majorly viable and showed ~19 % cell death (Figure 4i and Figure S7b), indicating it to be significantly less toxic than HMF. To probe the fibril-mediated cell death, we also analyzed the expression level of various genes involved in cell cycle and apoptosis by quantitative real-time polymerase chain reaction (qRT-PCR). Cells treated with fibrils showed increased expression of genes associated with apoptotic pathways in comparison to the buffer treated control cells (Figure 4j). Interestingly, the HMF displayed significantly higher expression for most of the apoptotic genes in contrast to the cells treated with PMF and comparable gene expression with Syn fibrils (except for p53, Bcl2 and Bcl-XL) (Figure 4j and k), further suggesting the toxic nature of HMF.

### Differential cellular uptake of α-Syn fibril polymorphs

Growing evidence suggests that α-Syn amyloids propagate in a prion-like manner and exhibits seeding and transmissibility properties^42^. Three distinct mechanisms govern its prion-like behavior: (i) the uptake of α-Syn seeds by cells, (ii) the seeding and templating of endogenous α-Syn *in cells*, and (iii) the transmission of aggregated α-Syn from one cell to another^42,43^.

First, we studied the internalization ability of the fibril polymorphs with equal-sized seeds in neuronal cells using fluorescein isothiocyanate (FITC) labeled fibrils (10% labeled and 90% unlabeled protein). We exogenously added FITC-labeled fibril seeds (at 0.1, 0.5, and 1 μM concentration) to the culture medium of SH-SY5Y neuroblastoma cells and analyzed after 24 h using confocal microscopy (Figure 5a). For all the fibril polymorphs, we observed a direct correlation in the fibril uptake and seed concentration. However, HMF and Syn fibrils displayed slightly higher internalization ability even at low seed concentrations compared to PMF (Figure 5b). Further quantification by FACS analysis confirmed a concentration-dependent increase in the number of FITC^+^ cells for all the fibril types (Figure 5c). The number of FITC^+^ cells in case of Syn fibrils and HMF increased from 26 % (0.1 μM) to 96 % (0.5 μM) and 10 % (0.1 μM) to 86 % (0.5 μM), respectively, suggesting nearly similar internalization ability of both the fibril types. PMF, however, displayed negligible internalization (~2 % FITC^+^ cells) at 0.1 μM, which increased to 77 % with the increase in the seed concentration (0.5 μM). For absolute quantification of the amount of fluorophore inside the cell, we compared the mean fluorescence intensity (MFI) values. As evident from the correlation plot in Figure 5d, the cells treated with PMF mostly showed FITC^+^ signal at higher concentrations, but the MFI values (705 and 1448) were two-fold lower than HMF (1443 and 2837) and Syn fibrils (1305 and 2738) at 0.5 and 1 μM seed concentration, respectively (Figure 5d). This difference in the MFI values clearly indicates that although 80-90 % cells are positive for FITC in the presence of all fibril seeds at higher concentrations (≥ 0.5 μM), the amount of fibrils internalized in neuronal cells varies significantly between the two polymorphs (PMF and HMF), with higher uptake efficiency observed in case of HMF.

**Figure 5:**
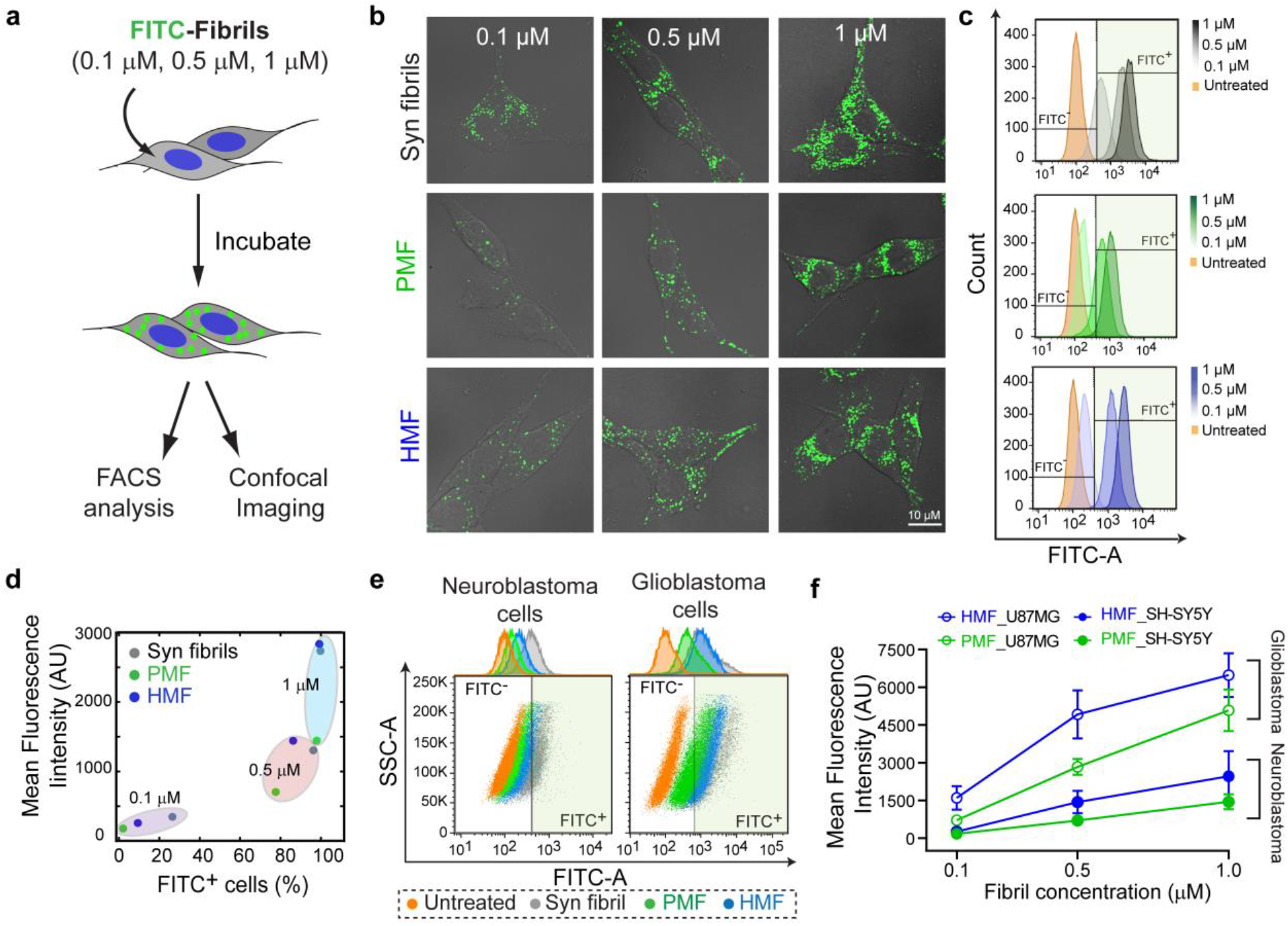
Internalization ability of α-Syn fibril polymorphs *in cells*. (a) Schematic representation of the experimental design for internalization of FITC-labeled fibril seeds at different concentrations and the formation of punctate-like structures *in cells*. (b) Confocal images of SH-SY5Y cells treated with FITC-labeled Syn fibrils, PMF and HMF seeds at three different protein concentrations (0.1 μM, 0.5 μM, 1 μM). Scale bar is 10 μM. (c) Histogram plots depicting a concentration-dependent increase in the FITC signal upon internalization of fibril polymorphs (PMF and HMF) along with Syn fibrils. The gating for FITC^−^ (non-shaded area) and FITC^+^ (shaded area) is based on the fluorescence intensity of the untreated sample. (d) The correlation plot of mean fluorescence intensity (Y-axis) and percentage cells positive for FITC (X-axis). The MFI values for each fibril sample are encircled (grouped) according to their concentration. Although 80-90 % of cells treated with PMF ≥ 0.5 μM concentration are FITC^+^, the amount of fibrils internalized (indicated by respective MFI values) is less than HMF and Syn fibrils. (e) The overlaid dot-plot of SH-SY5Y (neuroblastoma) and U87-MG (glioblastoma) cells treated with 0.1 μM fibril samples analyzed by FACS. The gating for FITC^−^ (non-shaded area) and FITC^+^ (shaded area) is based on the fluorescence intensity of the untreated sample in each case. (f) Comparative mean fluorescence intensity plot of neuroblastoma and glioblastoma cells treated with the HMF and PMF at 0.1 μM, 0.5 μM, and 1 μM concentrations. The data shows higher uptake efficiency of fibrils in glioblastoma compared to neuroblastoma cells. The values in (d) and (f) represent mean and mean ± s.e.m, respectively with n=3 from independent experiments.

Recently, it has been demonstrated that different α-Syn fibril strains target distinct cell types and regions within the brain, accounting for differences in the pathology observed among synucleinopathies^18^. Therefore, to examine whether the internalization ability of the polymorphs is altered upon changing the cell type, we performed the internalization assay in a glioblastoma cell line (U87-MG) under identical conditions. We specifically used the glial cell type, as α-Syn aggregates are known to form glial cytoplasmic inclusions in MSA^44^. Similar to neuronal cells, an increase in the fibril uptake in U87-MG cells was also observed with an increase in the seed concentration for all the fibril types (Figure S8a). The polymorphs, however, showed significant differences in the amount of fibrils internalized. There was high uptake of Syn fibrils (1480 MFI) and HMF (1409 MFI) even at low (0.1 μM) seed concentration with ~75 % and ~68 % FITC^+^ cells, respectively (Figure S8b and c). A subsequent increase in the Syn fibrils and HMF uptake was observed at seed concentration ≥ 0.5 μM with ~99 % FITC^+^ cells. On the contrary, only ~36 % of cells treated with PMF showed a positive signal for FITC at 0.1 μM seed concentration with 714 MFI (Figure S8b and c). With the increase in the concentration of PMF, the number of FITC^+^ cells increased, but the amount of fibril uptake remained low compared to Syn fibrils and HMF, as indicated from the correlation plot (Figure S8c). Taken together, the internalization pattern amongst polymorphs remains the same for the two cell lines. However, the extent of fibril uptake is greater for glioblastoma cells compared to neuroblastoma cells (Figure 5e and f). This difference is also apparent from the confocal images of neuroblastoma and glioblastoma cells treated with fibril polymorphs and their respective MFI values (Figure 5b, S8a and 5f).

### Intracellular seeding and aggresome formation by α-Syn polymorphs

One of the essential characteristics in the prion-like propagation of amyloid fibrils is the seeding and templating ability of the pathogenic protein aggregates^45^. To test whether PMF and HMF polymorphs possess different seeding potency *in cells*, we used SH-SY5Y cells stably expressing FLAG and tetracysteine-tagged α-Syn (C4-α-Syn). It is a robust cell line, and exhibits controlled expression of α-Syn upon doxycycline (DOX) induction. The small tetracysteine (C4)-tag (~1.5 kDa) binds to biarsenical dyes like FlAsH (green fluorescence) and ReAsH (red fluorescence) enables to study protein interactions and localization *in cells*^46–48^. C4-α-Syn exhibits identical biophysical and biochemical characteristics as that of wild-type α-Syn^49^, suggesting it to be a suitable model system for studying in cell seeding and aggregation. Post 24 h treatment with fibril seeds (Figure 6a), we observed (i) the formation of punctate structures (green; G) (Figure 6b, magnified box 2 and 4) arising from the aggregation of endogenous α-Syn, and (ii) the colocalization of rhodamine(Rh)-labeled fibril seed with endogenous protein (yellow; Y) (Figure 6b, magnified box 1 and 3), demonstrating the seeding event. The majority of the cells treated with Syn fibrils and PMF showed the formation of endogenous punctate-like structures (Figure 6b, Syn fibrils and PMF), along with colocalization with Rh-labeled internalized fibril seeds. However, the cells treated with HMF displayed mostly diffused expression of C4-α-Syn with low numbers of punctate structures (Figure 6b, HMF). Given that the amount of the internalized fibril differs amongst the fibril polymorphs (at 0.1 μM concentration), we assessed the seeding and fibril amplification of endogenous α-Syn with respect to the levels of internalized fibrils (Figure 6b and c). We quantified the total number of endogenous punctate structures (green, G) and total internalized fibril seeds (red, R) in each case and calculated the G/R ratio. PMF displayed nearly 1.5-fold (G/R) increase in the amplification of endogenous protein with respect to the internalized fibrils, whereas it was significantly low (~0.3-fold) in case of HMF (Figure 6c). Interestingly, even the low amounts of PMF seeds were sufficient enough to seed and amplify the endogenous protein, which was also higher than the fold increase (i.e. 0.85-fold) observed for α-Syn fibrils (Figure 6c). This was further evident from the percentage of relative seeding event (Y/R %), which was significantly higher in cells treated with PMF (43 %) and Syn fibrils (35.6 %) compared to HMF (9.8 %) (Figure 6d).

**Figure 6:**
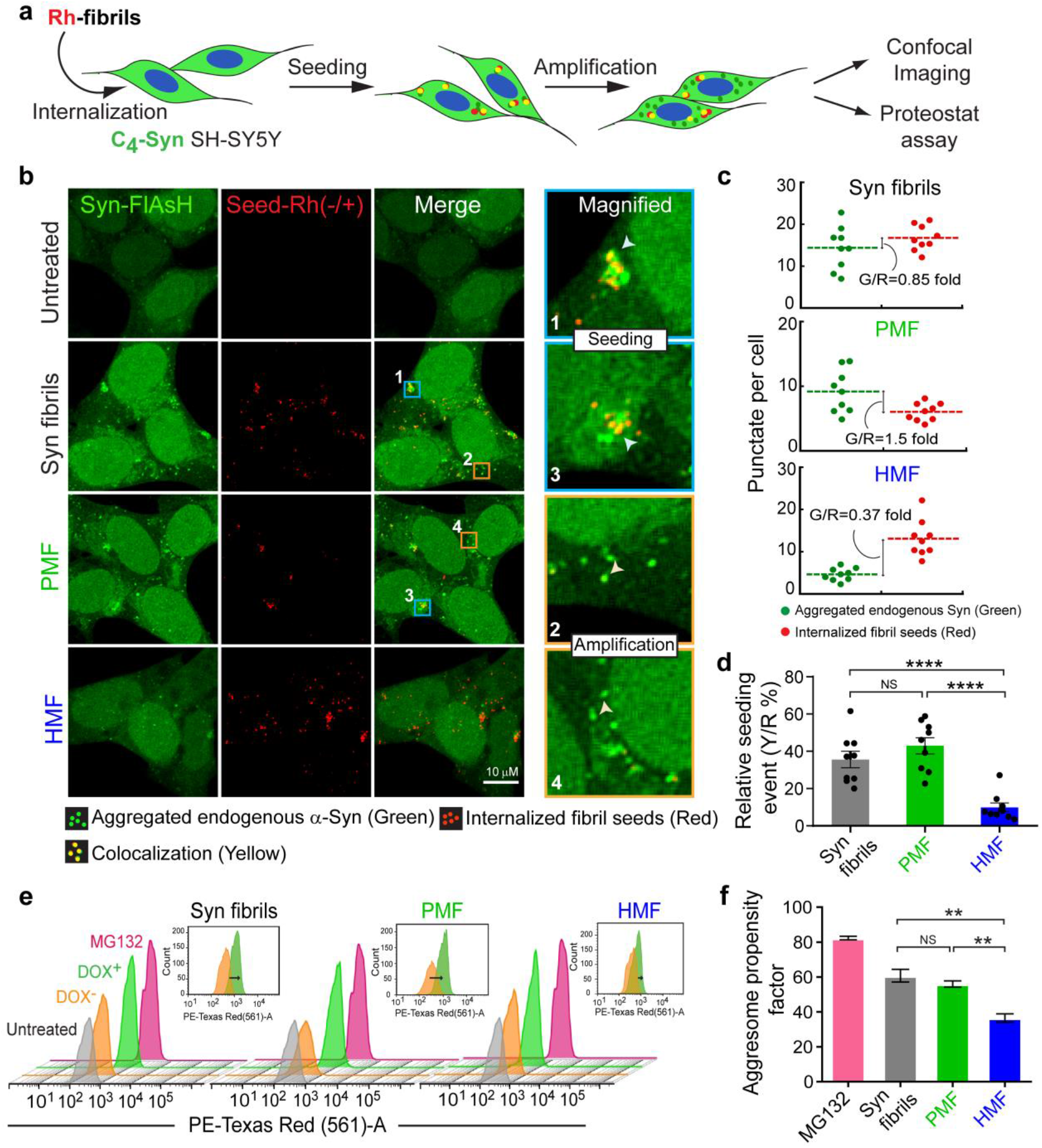
*In cell* seeding of α-Syn fibril polymorphs. (a) Schematic of experimental design to study the seeding and amplification of endogenous protein (α-Syn) upon exogenous addition of rhodamine (Rh)-labeled fibril seeds. (b) Confocal images of the SH-SY5Y cells overexpressing C4-α-Syn treated with 100 nM of Rh-labeled Syn fibrils, PMF and HMF seeds for 24 h. Right panel shows magnified images demonstrating seeding (box 1, 3) and amplification (box 2, 4) of the endogenous protein by Syn fibrils and PMF. Scale bar is 10 μM. (c) Quantification of endogenous protein aggregates with respect to internalized fibril seed. The plot shows the number of punctate structures (green; G) and internalized fibril seed (red; R) per cell along with G/R ratio indicating the fold change in number of endogenous aggregates with respect to the internalized fibrils. The data is quantified from random images from two independent sets with n > 100 cells using ImageJ. (d) Quantification of relative seeding event (%) observed at 24 h post-treatment of cells with fibril seeds. The bar plot shows the values obtained by taking the ratio of number of observed seeding events (yellow; Y) and internalized fibril seed (red; R) per cell (Y/R) expressed in percentage. The data is quantified from random images from two independent sets with n > 100 cells using ImageJ. (e-f) Nature of endogenous aggregates determined using ProteoStat dye staining. (d) Overlaid staggered plot of fluorescence intensities (acquired in PE-Texas-Red A channel) of cells treated with Syn fibril polymorphs. Inset shows overlay of fluorescence intensity in DOX^−^ and DOX^+^ cells in Syn fibrils, PMF and HMF. The black arrow marks the shift in the fluorescence intensity upon binding of ProteoStat dye with aggresome in DOX^+^ cells. (b) Aggresome propensity factor from ProteoStat detection assay using FACS analysis (values represent mean ± s.e.m, n=3 from independent experiments). The statistical significance in (d) and (f) is determined by one-way ANOVA followed by Newman Keuls Multiple Comparison post hoc test; *P≤0.05; **P≤0.01; ***P≤0.001; **** P≤0.001 NS P>0.05.

Furthermore, to assess the nature of the aggregates formed by PMF and validate the seeding results, we performed quantitative flow cytometric analysis with the aggresome specific marker ProteoStat^50^. A clear shift in the mean fluorescence intensity from DOX^−^ to DOX^+^ cell population was observed in Syn fibrils and PMF, suggesting the formation of aggresomal inclusions after 48 h (Figure 6e, and inset). However, the shift in the fluorescence intensity was considerably less in the case of cells treated with HMF. The difference in the binding of ProteoStat dye with endogenous aggregates in the cells treated with HMF and PMF was also evident from the calculated aggresomal propensity factor, which was significantly low in cells treated with HMF (36 %) in comparison to PMF (56 %) and Syn fibrils (60 %) (Figure 6f). Overall, the data imply that even trace amounts of PMF seeds are sufficient enough to induce abundant α-Syn pathology and trigger the formation of aggresomes *in cells*.

### Cell-to-cell transmission of α-Syn polymorphs

In the past few years, the concept of spreading of misfolded protein aggregates from one region of the brain to another in a prion-like fashion has been proven in various neurodegenerative disorders^8^. To accomplish the spread of α-Syn pathology via cell-to-cell transmission, protein aggregates must travel from one cell to another cell to reach the potential sites. For studying cell-to-cell transmission ability of α-Syn polymorphs, we used FITC and Rh-labeled fibrils, each of which were allowed to internalize in two different sets of cells maintained under identical conditions (Figure 7a). After co-culturing the two different sets of cells (Figure 7a), we examined the trans-cellular spread of fluorescently labeled aggregates by confocal microscopy (Figure 7b). The data revealed ~50 % and ~43 % FITC^+^Rh^+^ cell population in case of Syn fibrils and HMF, respectively (Figure 7c), demonstrating the cell-to-cell transmission of these two fibril types (Figure 7b, white arrowhead). In contrast, PMF displayed comparatively low cell-to-cell transmission ability with only ~19 % FITC^+^Rh^+^ cells. Further, we validated the imaging results more quantitatively by flow cytometry (Figure 7d and e). FACS analysis revealed 45% FITC^+^Rh^+^ cells in case of Syn fibrils closely followed by HMF with 32 % FITC^+^Rh^+^ cells. Whereas, only 9 % cells were positive for both FITC and Rh upon treatment with PMF (Figure 7d and e). Therefore, HMF can travel from one cell to another, suggesting its ability to infect and spread α-Syn pathology like infectious prions, while PMF clearly lack these abilities.

**Figure 7:**
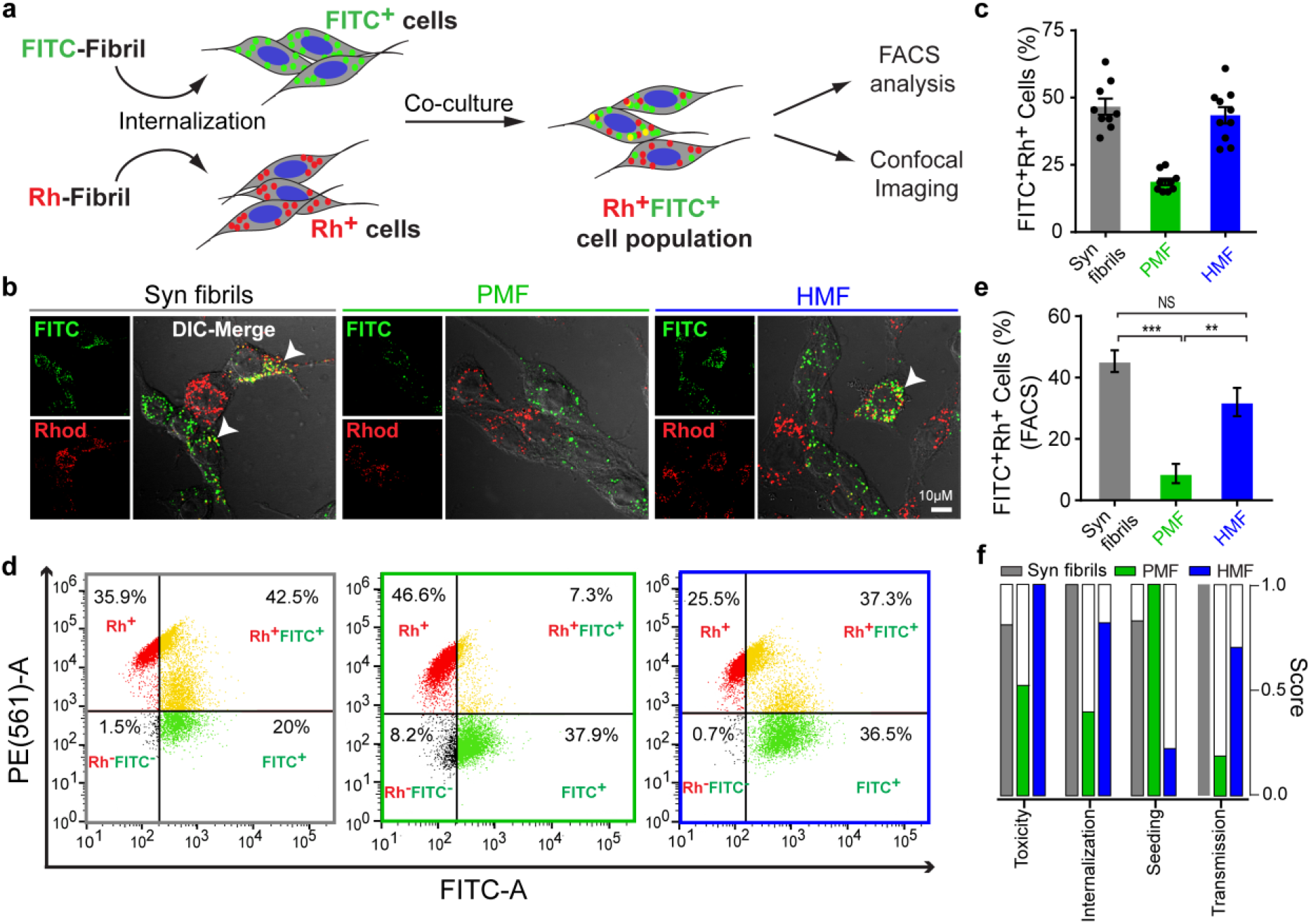
Cell-to-cell transmission of α-Syn fibril polymorphs. (a) Schematic representation of the experimental design to study cell-to-cell transmission of fibrils. FITC- and Rh-labeled fibril seeds of each polymorph are internalized in two different sets of neuroblastoma cells, followed by co-culturing the FITC and Rh labeled cell population. (b) Confocal images of the SH-SY5Y cells showing co-cultured Rh^+^ and FITC^+^ cell population treated with Syn fibrils, PMF and HMF seeds for 24 h. Cells positive for both FITC and rhodamine in case of Syn fibrils and HMF are marked in white arrowhead. The scale bar is 10 μM. (c) Quantification of percentage population of FITC^+^ Rh^+^ cells calculated from confocal images. The data is quantified from n>200 cells using ImageJ. (d) Quadrant plot obtained from FACS analysis of co-cultured cells treated with fibril samples showing the percentage of cells in each subpopulation (Rh^−^ FITC^−^ (black); Rh^+^ only (Red); FITC^+^ only (green) and FITC^+^ Rh^+^ (yellow), n=3. (e) Quantification of percentage population of FITC^+^ Rh^+^ cells from FACS analysis. The values represent mean ± s.e.m, n=3 from independent experiments. The statistical significance is determined by one-way ANOVA followed by Newman Keuls Multiple Comparison post hoc test; **P≤0.01; ***P≤0.001; NS P>0.05. (f) Comparative plot of toxicity and prion-like properties exhibited by Syn fibrils, PMF and HMF. Scoring is based on the experimental data sets. Scores are normalized with respect to the maximum value (0,1). The data suggests that more structured polymorph (HMF) is potentially more pathogenic compared to less structured PMF.

## Discussion

In synucleinopathies, the involvement of α-Syn aggregation in diseases with distinct clinicopathological conditions is not fully understood yet. In this context, recent studies have reported the generation of α-Syn fibril polymorphs that possess distinct structural and biochemical properties under different experimental conditions^13,15,16,19,51^. These findings indicate that the same precursor protein can form amyloids with different structures leading to distinct pathologies. The aggregation of α-Syn involves the formation of an aggregation-competent nucleus^52,53^, which is a stochastic process and may not occur in a controlled and definite manner *in vivo*^54^. As a consequence, the intermediate species formed during the aggregation cascade may result selectively into one or the other type of structural assemblies depending upon environmental and cellular conditions. We hypothesized that the heterogeneous species formed during the aggregation pathway result in fibril polymorphism. In this study, we developed two α-Syn polymorphs i.e., PMF and HMF, from the intermediates formed during the nucleation and early elongation phase of α-Syn aggregation. Both these polymorphs differ in their secondary structural components and fibrillar alignment/packing, as observed by FTIR and X-ray diffraction studies (Figure 1). Both the polymorphs also showed different proteinase K-digestion profiles, suggesting distinct structural arrangements as well as compactness in the fibril core structure. Indeed, the HDX-MS and ssNMR studies highlight the internal structural variations in the fibril core of both the fibril polymorphs. For example, PMF is shown to be more prone to amide exchange than HMF, indicating this polymorph to be less ordered (Figure 2). The ssNMR further supports residue-level structural differences, particularly in their NAC domain (65-80 region), suggesting the NAC region of PMF is more flexible compared to HMF (Figure 3). In this context, previous studies showed α-Syn in LBs and GCIs from PD and MSA patients, respectively, are structurally different and show the presence of diverse proteolytic fragments upon limited proteolysis^9^.

One of the important property which emerges due to the structural differences among polymorphs is their different templating/seeding capability^15,55^. A recent study by Soto and co-workers also provides direct evidence of differential seed-mediated protein amplification from cerebrospinal fluid samples of PD and MSA patients^56^. When we analyzed the seeding efficiency of the polymorphs, we found that PMF displays significantly higher nucleation and seeding potential compared to HMF (Figure 4), supporting that internal structure and/or fibril packing influences the seeding capacity of these polymorphs. Not only the seeding, it also influences the surface hydrophobicity of the polymorphs, which in turn results in distinct levels of cytotoxicity (Figure 4).

α-Syn amyloids propagate from one region of the brain to another through cell-to-cell transmission and seeding in various stages of PD pathology^8,57^. We hypothesized that α-Syn polymorphs exhibit distinct seeding and transmission ability *in cells*, reminiscent of prion strains. HMF readily internalizes in both neuronal and glial cells and transmits from one cell to another, whereas PMF displays comparatively low uptake in these cell types and lacks the transcellular spreading ability (Figure 5 and 7). Interestingly PMF, however, exhibits high seeding potency, both *in vitro* and *in cells* (Figure 4 and 6). These findings corroborate well with a recent study^14^, which indicates that the internalization ability (i.e., a high number of internalized seeds) of polymorph might not correlate with its templating/seeding potency. Rather, seeding ability would highly depend on the capability of the seeds to carry out the complementary interactions with the soluble monomeric species and, therefore, the rate of recruitment of endogenous monomers. The strain-specific seeding might also be governed by seed conformation and cellular milieu^9,58^. This has been particularly shown with brain extracts from MSA or PD patients, which also exhibit distinct seeding potency *in vitro*^58,59^ and *in vivo*^9^.

The present data, therefore, suggest that conformational heterogeneity in the aggregation pathway may result in the formation of fibril polymorphs with different structures, toxicity and prion-like behavior *in* cells (Figure 7f and S8). Based on different cellular conditions, the population and/or lifetime of these intermediates may dictate the overall population of one polymorph over the other, which eventually imparts the consequent pathology. The present study would significantly help to understand the complexity of the α-Syn aggregation pathway and resultant polymorphs associated with PD and related disorders.

## Materials and Methods

### Chemical and Reagents

All chemicals used in the study were of the highest purity available and obtained from Sigma Chemical Co. (St. Louis, MO, USA) and HiMedia (India), unless otherwise specified. ^13^C-glucose and ^15^Nlabeled ammonium chloride were purchased from Cambridge isotope laboratories, Inc. (USA). Cell culture media (DMEM) and fetal bovine serum (FBS) were obtained from GIBCO (USA), MEM (supplemented with non-essential amino acids) from HiMedia (India) and Tet system approved FBS was obtained from Takara Bio USA. FITC and rhodamine dyes were purchased from Thermo Fisher Scientific and FlAsH-EDT2 from Cayman Chemicals (USA). The kits used in the study were: FITC Annexin V apoptosis detection kit (BD Biosciences, USA), ProtoScript-II first-strand cDNA synthesis kit (NEB, USA), Brilliant-III ultrafast SYBR Green qPCR master mix (Agilent Technologies, Santa Clara, CA, USA) and ProteoStat-aggresome detection kit (Enzo Life sciences, USA). Milli-Q system (Millipore Corp., Bedford, MA) was used for double distillation and deionization of water.

### Protein expression and purification

Expression and purification of recombinant α-Syn was done using pRK172 plasmid as per the protocol described by Volles *et al*^60^ with minor modifications^61^. The purity of the protein was assessed by SDS-PAGE and MALDI-TOF mass spectrometry. Solid, lyophilized α-Syn was dissolved in 20 mM glycine-NaOH buffer (pH 7.4, 0.01% NaN3) with the desired concentration. LMW α-Syn, which mostly consists of monomeric protein was used throughout the aggregation study and prepared as per previously published protocol^23^.

### Aggregation kinetics study and isolation of different α-Syn fibril variants

300 μM of LMW protein prepared in 20 mM glycine-NaOH, pH 7.4 containing 0.01% NaN_3_. was incubated at 37 °C in Echo Therm model RT11 rotating mixture (Torrey Pines Scientific, USA) at 50 rpm. Aggregation kinetics was continuously monitored by measuring CD and ThT fluorescence at regular intervals. After incubation of LMW α-Syn and its subsequent conversion into fibrils (confirmed by β-sheet structure (minima at 218 nm) in CD study and high ThT binding), the solution was ultra-centrifuged at 30,000 rpm to obtain a pure fibril population. The concentration of the fibril pellet was calculated by measuring the concentration of supernatant by taking absorbance at 280 nm and subtracting it from the initial concentration (300μM). Finally, the fibril pellet was re-dissolved into buffer to obtain the desired concentration and was referred to as Syn fibrils.

During aggregation kinetics, when helix-rich conformation was detected in CD, α-Syn aggregation mixture was centrifuged (14,000 g, 20 mins at 4 °C) and a fibril pellet was obtained, which consisted of pre-matured fibrils, named as PMF. The concentration of the fibril pellet was calculated by subtracting the concentration of supernatant (A_280_ nm) from the initial concentration of protein solution (i.e. 300 μM) as described previously^23^. The supernatant obtained after pelleting down the pre-matured fibrils was collected in a fresh microfuge tube and its concentration was determined (by measuring A_280_ nm). It was then was passed through 100 kDa cut off filter (Centricon YM-100, Millipore) and retentate (containing helix-rich oligomers) was transferred in a fresh tube. The concentration of the retentate was calculated by subtracting the concentration (A_280_ nm) of flow-through from the supernatant concentration. Thereafter, the isolated helix was immediately incubated at 37 °C for another 8 days and allowed to mature into fibrils. After incubation, the resulting fibrils were ultra-centrifuged at 30,000 r.pm. to obtain pure fibril population and its concentration was determined as done for Syn fibrils. The fibril pellet was re-dissolved into buffer to obtain desired protein concentration and labeled as HMF.

### Circular dichroism (CD) spectroscopy

The fibril samples were prepared as described above. CD measurements of the fibril samples diluted in 20 mM glycine-NaOH buffer (pH 7.4,0.01% sodium azide) at final protein concentration of 7.5 μM (200 μl) were performed using JASCO-1500 CD spectrophotometer as per the previously published protocol^23^. The experiment was repeated thrice with independent samples.

### ThioflavinT (ThT) fluorescence assay

For ThT fluorescence assay, each fibril sample was diluted in 20 mM glycine-NaOH buffer (pH 7.4), 0.01% sodium azide at a final concentration of 7.5 μM (200 μl) and 2 μl of 1mM of ThT dye prepared in Tris-HCl buffer (pH 8.0, 0.01% Sodium azide) was added to it. The assay was performed using FluoroMax-4 spectrofluorometer (HORIBA Jobin Yvon) instrument using the settings as previously reported^23^. Three independent sets of experiments were done.

### Fourier-transform infrared (FTIR) spectroscopy

10 μl of the α-Syn fibril samples were spotted on a thin KBr pellet and allowed to dry. FTIR was performed using BrukerVertex-80 instrument equipped with a DTGS detector using the parameters as described previously^62^. The deconvolution was carried out from the raw data corresponding to the amide-I region (1700-1600 cm^−1^) using by Fourier self-deconvolution (FSD) method. The deconvoluted spectra were put through Lorentzian curve fitting procedure and integrated using opus-65 software as per manufacturer’s instructions. Three independent recordings were done from three different fibril preparations.

### Atomic force microscopy (AFM)

Diluted fibril samples (50 μM in 20 mM glycine-NaOH buffer, pH 7.4, 0.01% sodium azide) were spotted on a small freshly cleaved mica sheet and incubated at room temperature (RT) for 5-7 mins. It was washed with milli-Q water to remove any unbound protein and dried in a vacuum desiccator for 30 mins at RT. Images were taken in tapping mode with a silicon nitride cantilever using Veeco Nanoscope IV Multimode AFM with 1.0 Hz scan speed. Three to four different areas from two independent samples were imaged.

### Transmission electron microscopy (TEM)

For TEM sample preparation, 10 μl of fibril sample (50 μM) was spotted on carbon-coated Formvar EM grids (Electron Microscopy Sciences, PA, USA), incubated for 5 mins and stained with uranyl formate (1% v/v). Imaging was done using transmission electron microscope (Philips CM-200) with a magnification range of 6,600 X to 12,000X at 200 kV. Recording of images was done digitally with the aid of Keen View Soft imaging system. Random 10-12 images were acquired for each sample.

### X-ray diffraction pattern of fibrils

For X-ray diffraction study, samples were loaded into a clean pre-dried 0.7 mm capillary tube and dried overnight under vacuum. The capillary tube with dried protein sample was mounted in the path of X-ray beam at 1.2 kW for 300 sec. The images were obtained using Rigaku R Axis IV++ detector (Rigaku, Japan). Finally, the diffraction data were analyzed using Adxv software. Two to three diffraction patterns were collected from at least two independent samples.

### Proteinase K digestion assay

Polymorphs along with Syn fibrils at the protein concentration of 100 μM dissolved in 20 mM glycine-NaOH, pH 7.4, 0.01% NaN3 were treated with Proteinase K (10 μg/ml) at 37 °C and aliquots were collected at given intervals. The treatment was done in two sets; one for (i) SDS-PAGE and another for (ii) MALDI-TOF mass spectrometry as described below.

At each time point, sample buffer (50 mM Tris-HCl, pH 6.8, 4% SDS, 2% β-mercaptoethanol, 12% glycerol and 0.01% bromophenol blue) was added in the aliquots and the reaction was stopped by heating the samples at 95 °C on dry bath for 5 mins. 15 μl of samples were loaded onto 18% Tris-tricine SDS-PAGE and the digestion profile of different fibrils was analyzed. For MALDI-TOF, the digested fibril samples were mixed with sinapinic acid (trans-3,5-dimethoxy-4-hydroxycinnamic acid; SA matrix) in the ratio 1:1 (2 μl sample + 2 μl SA). The mixture was immediately spotted on the target plate and allowed to dry at RT for 10-15 mins. MALDI-TOF/TOF mass spectrometry was performed as previously described^63^. Three independent spectra for each sample were recorded for individual time points.

### Hydrogen/ deuterium (H/D) exchange-Mass spectrometry (MS) measurements

#### Peptide mapping

A peptide map of α-Syn digestion was generated as described previously^31^. Briefly, the protein was dissolved in water before subjecting to on-line pepsin digestion using an immobilized pepsin cartridge (Applied Biosystems, USA). The eluted peptides were collected using a peptide trap column and eluted further from the column using an acetonitrile gradient. The peptides were subjected to the Synapt G2 mass spectrometer (Waters) for sequencing and analyzed using the Protein Lynx Global Server software (Waters) and manual inspection.

#### HDX-MS measurements

Fibril samples were re-suspended in 20 μl of 20 mM glycine-NaOH buffer to get the final 300 μM concentration. 10 μl of the sample was diluted into 190 μl of the same buffer (without EDTA) prepared in D_2_O and labeling pulse was given for 30 s, 1800 s and 18000 s pulse (in deuterated buffer at pH 8.0, 37 °C). After the pulse, 400 μl of ice-cold quench buffer (0.1 M glycine-HCl, 8.4 M Gdn HCl, pH 2.5) was added to the sample and incubated on ice for 1 min to dissolve the fibrils. The above sample was desalted using a HiTrap desalting column equilibrated with pH 2.5 water using the Akta chromatography system. Thereafter, these desalted samples were injected into a nano ACQUITY UPLC system (Waters Corp, USA) using an immobilized pepsin cartridge for online pepsin digestion. Further processing was carried out using a Waters Synapt G2 mass spectrometer as described previously^64^. The percentage deuterium incorporation (%) for peptides displaying a bimodal mass distribution was calculated^64,65^ and heat map was plotted using GraphPad. The bimodal mass distributions were then fitted to the sum of two Gaussian distributions using OriginPro 8 (Origin Lab, USA) to obtain centroid mass for each peak. The experiment was repeated thrice from three independent fibril preparations.

### Solid-State Nuclear Magnetic Resonance (ssNMR) spectroscopy

Uniformly labeled- ^15^N, ^13^C α-Syn protein was expressed in *E. coli* BL21 (DE3) strain in M9 medium containing ^13^C-glucose and ^15^N NH_4_Cl as the sole source of carbon and nitrogen, respectively. Thereafter, the protein was purified in the same way as for unlabeled protein (mentioned earlier). In order to obtain high concentration of ^15^N, ^13^C-labeled fibril polymorphs for ssNMR, the aggregation reaction of 300 μM of LMW ^15^N,^13^C-labeled α-Syn protein in ~8-10 ml of 20 mM glycine-NaOH (pH 7.4) buffer was carried out and the fibril polymorphs were prepared similarly as mentioned earlier.

All the samples were ultra-centrifuged at 200,000xg for 1 h before filling into the rotor. The sample was filled in 4 mm Bruker rotor using the swinging bucket centrifuge at 4000xg. Trace amount of DSS was added into the rotor to use it as internal reference. The sample temperature was maintained around 10 °C for all the measurements. The experiments were done using Bruker 700 MHz Avance-III for sequential assignment of fibrils sample. These experiments (DARR, NCA, NCACB, NCACX, CANCO and NCOCX) were done as described previously^33^. The detailed experimental parameters are listed in Table S1.

The assignments were performed by the standard backbone walk approach using multiple 3D spectra and relying on the redundancy of chemical shifts. The residues were assigned by starting a walk form threonine or serine residues. These residues have distinct Cα and Cβ chemical shifts and can be easily identified in different spectra. In situation when the walk could not be carried out, the next alanine and threonine peaks were chosen to start a fresh walk. The pairs of residues identified in walk were assigned based on the uniqueness of these pairs in the sequence. The repeating pairs of amino acids were assigned if only one such pair was unassigned in the sequence. There are few peaks in the Syn fibril that were assigned based on largely similar HMF assignment, especially in the 1-40 and 80-98 regions.

### Optimization of fibril seed length

All the fibril samples were sonicated using probe sonicator (Sonics & Materials Inc., Newtown, CT, USA) at 20 % amplitude, 3 sec on-1 sec off pulse for 1 min, 2 min and 3 min. For each time point, TEM analysis was done as described earlier. The fibril length was calculated from different fields with >270 counts using Image J software.

### Aggregation kinetics in the presence of seeds

Pre-formed fibril seeds (0.1% v/v) were incubated with freshly prepared LMW (150 μM) at 37 °C with slight shaking in 96-well clear bottom plate. Thioflavin T was added along with the samples in a 96-well plate to monitor the aggregation kinetics. LMW, fibrils seeds, buffer and ThT alone were incubated under identical conditions as controls. An increase in ThT fluorescence signal (excitation at 450 nm) was measured at regular intervals using SpectraMax M2e microplate reader (Molecular Devices, USA). The obtained data were fitted in a sigmoidal growth curve using OriginPro 8.0 (Origin Lab, USA). The experiment was repeated thrice.

First derivative and second derivative plots were obtained from each of the fitted curves of the aggregation kinetics. First maxima and final minima were obtained from the second derivative plots. The inverse of the time taken to reach the first maxima of the second derivative plots correspond to the nucleation rate of the aggregation kinetics, respectively.

The lag time was calculated according to the published protocol^38^ using the following equation

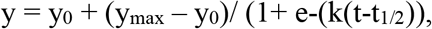

where y is the ThT fluorescence at a particular time point,

y_max_ is the maximum ThT fluorescence and y_0_ is the ThT fluorescence at t_0_ and t_lag_ was defined as

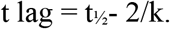

### ANS Binding

ANS binding assay was performed as previously described^23^. Briefly, 3 μl of 5 mM ANS was added to diluted fibril samples (7.5 μM, 200 μl) and incubated for 10 mins in dark at RT. The assay was performed using FluoroMax-4 spectrofluorometer (HORIBA Jobin Yvon) with 370 nm as an excitation wavelength and 400-600 nm as an emission wavelength range. The acquired spectra were plotted in OriginPro 8 (Origin Lab, USA). The experiment was repeated thrice.

### Nile Red binding assay

1 mM stock solution of NR dye was prepared in DMSO and 0.2 μl of 1 mM NR dye (from the prepared stock) was added to 200 μl of the 10 μM fibril samples followed by 5 min incubation at RT. Fluorescence measurements were taken at 550 nm (excitation wavelength) and 565-720 nm (emission wavelength range) using FluoroMax-4 spectrofluorometer (HORIBA Jobin Yvon). The acquired spectra were plotted in OriginPro 8 (Origin Lab, USA). The experiment was repeated thrice with independent fibril preparations.

### MTT assay

Human neuroblastoma SH-SY5Y cells (obtained from the National Centre for Cell Science, Pune, India) were cultured in DMEM (GIBCO) supplemented with 10 % FBS and 1% antibiotic (streptomycin and penicillin) at 37 °C in a humidified chamber with 5 % CO_2_. The cells were trypsinized and seeded in 96-well plate with 1×10^4^ cells per well and incubated overnight. Thereafter, cells were treated with α-Syn fibril polymorphs (25 μM) for 24 h. MTT reagent (0.5 mg/ml) was added to the cells followed by 4 h incubation. 100 μl solubilization buffer (DMF (50%), SDS (20%), pH 4.7) was added and plate was kept overnight at 37 °C. Absorbance at 560 nm and 690 nm (for background correction) was measured using SpectraMax M2e microplate reader (Molecular Devices, USA). Three independent experiments were performed.

### Annexin-V PI staining

For studying apoptosis, SH-SY5Y cells were seeded in 24-well plate at a cell density of 1×10^5^ cells per well as per culture conditions used for MTT assay. After 24 h, the cells were then treated with α-Syn fibril polymorphs (50 μM) and allowed to incubate for 48 h. Untreated and cells treated with buffer were kept as controls. Post-treatment, cells were stained with Annexin V-FITC and PI (Apoptosis Detection kit, BD Biosciences) as per manufacturer’s recommended protocol and were quantified in a flow cytometer (FACS Aria, BD Biosciences). The samples were analyzed using the BD FACS Diva software with 10,000 cells counted for each sample. The acquired data was plotted in FlowJo_V10 (Tree Star, Inc) software. Three independent experiments were performed for each set.

### Quantitative Real Time-PCR

SH-SH5Y cells seeded in a 12-well plate were treated with 50 μM of fibril samples for 48 hours. Untreated cells and buffer treated cells were kept as controls. Post incubation, the treated cells were trypsinized, pelleted and further subjected for RNA isolation using TriZol (Invitrogen) method as per manufacturer’s recommendation. The concentration of isolated RNA per μl was estimated using Nanodrop spectrophotometer. Thereafter, cDNA synthesis was carried out by ProtoScript-II first strand cDNA synthesis kit (NEB, USA) with random hexamer primers, in accordance with manufacturer’s protocol. SYBR Green method was used to perform quantitative Real-time PCR (qRT-PCR) with Brilliant-III ultrafast SYBR Green qPCR master mix (Agilent Technologies, Santa Clara, CA, USA) as per manufacturer’s protocol on an Agilent MX3000P. The primers GAPDH, p53, p21, BAX, Bcl-2, Bcl-XL, Cas3, Cas9, AIF and Bad were used for the respective genes (Table S2). The data was acquired from three independent experiments.

### Labeling of α-Syn monomer

LMW protein solution was prepared as described earlier. FITC and Rhodamine (Rh) labeling of LMW α-Syn was performed^62^ as per the manufacturer’s recommendations (Molecular Probes, USA). Post labeling of protein, 5 mg of lyophilized labeled α-Syn was dissolved in 500 μl of 20 mM glycine-NaOH buffer (pH 7.4) and its concentration was measured as per the manufacturer’s protocol (Molecular Probes). This was used as a FITC/Rh labeled protein stocks for further experiments.

For the setting up the aggregation and isolation of different labeled fibril samples, the unlabeled LMW α-Syn (90 %) was mixed with 10 % labeled α-Syn at a final protein concentration adjusted to 300 μM in 20 mM glycine-NaOH buffer (pH 7.4). This protein solution was incubated at 37 °C in rotating shaker (EchoTherm model RT11 (Torrey Pines Scientific, USA)) at 30 rpm until fibril formation and labeled fibril polymorphs were obtained as described earlier.

### Internalization of α-Syn fibril polymorphs

SH-SY5Y cells were cultured in DMEM (GIBCO) supplemented with 10 % FBS (GIBCO) and 1% antibiotic (streptomycin and penicillin) and seeded at the cell density of 1×10^4^ on glass coverslips in 24 well plates and 5×10^5^ cells directly in 12-well plate for 24 h at 37 °C with 5% CO_2_ in a humidified chamber. FITC-labeled fibril seeds were generated by sonication at 20 % amplitude with 3 sec on, 1sec off pulse for 3 mins (Syn fibrils) or 2 mins (PMF and HMF). The cells were then treated with different concentration of FITC-labeled fibril seeds (0.1 μM, 0.5 μM and 1 μM) for 24 h. Untreated and buffer treated cells were kept as controls. After 24 h incubation with samples, the cells were analyzed by flow cytometry (FACS Aria, BD Biosciences) in FITC channel. The gating was done based on the fluorescence intensity of the untreated sample. For each sample, 10,000 cells were analysed using the BD FACS Diva software. The acquired data was plotted and analyzed using FlowJo_V10 (Tree Star, Inc) software.

In the other set, cells seeded on the coverslips were fixed with 4% paraformaldehyde (PFA) post fibril treatment and washed twice with phosphate-buffered saline (pH 7.4). Coverslips were mounted on glass slides and imaged by Zeiss Axio-Observer Z1 laser scanning confocal microscope equipped with iPlan-Apochromat 63X/1.4 NA oil immersion objective. The experiment was repeated thrice. Image processing and quantification were done using Image J software.

U87-MG cells (obtained from the National Centre for Cell Science, Pune, India) were cultured in MEM (HiMedia) containing non-essential amino acids (NEAA) supplemented with 10 % FBS (GIBCO) and 1% antibiotic (streptomycin and penicillin). Internalization assay with human glioblastoma cell line U87-MG was performed using a similar protocol as mentioned above.

### *In cell* seeding experiment

Doxycycline inducible SH-SY5Y cells stably expressing C4-α-Syn^48,66^ were used for in *cell* seeding experiments. The stable SH-SY5Y cells were cultured at 37 °C in 5 % CO_2_ in DMEM (GIBCO) supplemented with 10 % Tet system approved FBS (Takara Bio USA) and 1% antibiotic. 10 ng of doxycycline hyclate (DOX) (Sigma, USA) was added for inducing the expression of C4-α-Syn. After 24 h of expression, the DOX-containing medium was replaced with medium containing 0.1 μM of Rh-labeled fibril seeds of polymorphs. Post 24 h treatment, the cells were stained with FlAsH-EDT2 using the previously described method^49^. Briefly, cells were incubated with 1 μM FlAsH dye (Cayman Chemicals, USA) and 10 mM EDT prepared in Opti-MEM (Invitrogen, USA), for 45 min at 37 °C incubator with 5 % CO2. Thereafter, the cells were washed with 100 mM EDT (prepared in Opti-MEM) for 10 min to remove any unbound dye. Cells were then fixed with 4% PFA, washed with PBS and mounted on the glass slides. Cell imaging was done using Zeiss Axio-Observer Z1 laser scanning confocal microscope (inverted) equipped with iPlan-Apochromat 63X/1.4 NA oil immersion objective. Image processing and the quantification of punctate (red; green and yellow) was done using cell counter plugin in ImageJ. N >100 cells were counted for each sample from independent sets of experiments.

### ProteoStat binding assay and flow cytometry analysis

For ProteoStat binding assay, we performed the experiment in two parallel sets; in (set I) the expression of C4-α-Syn was induced in cells by addition of 10 ng DOX followed by addition of unlabeled-fibril seeds (0.1 μM) for 24 h, marked as “DOX^+^ samples” and in (set II), cells were treated with unlabeled fibril seeds without any expression of C4-α-Syn, marked as “DOX^−^ samples”, used as controls. Cells treated with 5 μM MG132, a proteasomal inhibitor, were used as a positive control. Post 48 h of incubation, cells were fixed and subsequently stained using ProteoStat dye as per manufacturer’s recommendation (Enzo Life Sciences, USA). For FACS analysis, 1 × 10^6^ cells were fixed with 4 % PFA and fixed cells were permeabilized with 0.2 % Triton X-100. The working stock of ProteoStat was made by adding 0.5 μl of ProteoStat (5 μM in 5 ml of assay buffer) and added to the cells just before analysis. The cells were analyzed using a BD FACS Aria flow cytometer in Texas Red channel (BD Biosciences, CA, USA). All of the experiments were performed at least three times. Aggresome propensity factor (APF) was calculated by comparing the mean fluorescence intensity, given as APF = 100 × ((MFI_DOX_^+^–MFI_DOX_^−^)/MFI_DOX_^+^), where MFI_DOX_^+^ and MFI _DOX_^−^ are the mean fluorescence intensity (MFI) values from DOX^+^ cells and DOX^−^ cells treated with fibril samples. In this case, we used MFI of DOX^−^ cells as a control to subtract any fluorescence arising due to the binding of ProteoStat with internalized fibrils. The acquired data was plotted and analyzed in FlowJo_V10 (Tree Star, Inc) software.

### Cell-to-cell transmission

FITC and Rh labeled fibrils were prepared as described earlier. The labeled fibril seeds at the concentration of 1 μM were allowed to internalize in SH-SY5Y cells for 24 h. Post internalization, the cells were trypsinized and the equal number of cells treated with FITC and Rh-labeled fibril seeds were co-cultured and seeded in 6-well plate. Cells with single fluorophores were kept simultaneously as controls for 24 h at 37 °C in humidified CO_2_ chamber. Thereafter, cells were fixed with 4% PFA, washed with PBS and mounted on a glass slide. Cell imaging was carried out with Zeiss Axio-Observer Z1 laser scanning confocal microscope for identifying double-positive cells. The FITC^+^Rh^+^ cells from the acquired images were counted using cell counter plugin in ImageJ software with n >200 cells for each sample.

In another set, the co-cultured cells were trypsinized and analyzed by Flow cytometry in FITC and Rh channel using a BD FACS Aria flow cytometer (BD Biosciences, CA, USA) for quantification. The gating was done based on the fluorescence intensity of untreated samples and FITC^+^ and Rh^+^ alone cell populations. The experiment was repeated at least three times. The data acquired were analyzed and plotted in FlowJo V10 (Tree Star, Inc).

### Statistical Significance

The statistical significance was determined by one-way ANOVA followed by Newman Keuls Multiple Comparison post hoc test; *P≤0.05; **P≤0.01; ***P≤0.001; ****≤0.0001; NS P>0.05.

## Supplementary information

Supplementary information includes supplemental results, nine figures, and two tables.

## Author contributions

S.K.M and S.M. conceived the project and designed the experiments. S.M. performed all the experiments, unless otherwise noted. S.A with help of V.A carried out the solid-state NMR and H.K. carried out the HDX-MS study under the guidance of J.B.U. N.S. designed the primers for PCR and helped in the cell experiments. A.N. carried out flow cytometry and qRT-PCR. K.P performed the confocal imaging. P.K. and R.K performed electron microscopy imaging and N.N.J carried out FTIR. S.M and S.K.M wrote the manuscript. N.S. and A.N. helped in editing the draft. All the authors proof-read and approved the manuscript.

## Acknowledgment

We acknowledge Dr. Juan Gerez and Prof. Roland Reik, ETH Zurich, CH-8093 Zurich, Switzerland for the kind gift of C4-Syn SH-SY5Y cell line. We also acknowledge the Department of Biotechnology (DBT) – Basic Sciences, India [BT/PR22749/BRB/10/1576/2016], Government of India for the financial support. We are also thankful to IRCC and CRNTS, IIT Bombay for CM-TEM 200, FTIR, protein crystallography and Department of Biosciences and Bioengineering for FACS, confocal microscope, Bio-AFM, MALDI-TOF/TOF mass spectrometer facilities. S.M. acknowledges UGC (Govt. of India) for her fellowship.

## References

1 Goedert, M. α-synuclein and neurodegenerative diseases. Nat. Rev. Neurosci. 2, 492–501 (2001).

2 Jucker, M. & Walker, L. C. Self-propagation of pathogenic protein aggregates in neurodegenerative diseases. Nature 501, 45–51 (2013).

3 Goedert, M., Clavaguera, F. & Tolnay, M. The propagation of prion-like protein inclusions in neurodegenerative diseases. Trends Neurosci. 33, 317–325 (2010).

4 Luk, K. C. et al. Exogenous α-synuclein fibrils seed the formation of Lewy body-like intracellular inclusions in cultured cells. Proc. Natl. Acad. Sci. U. S. A. 106 (2009).

5 Volpicelli-Daley, L. A. et al. Exogenous α-synuclein fibrils induce Lewy body pathology leading to synaptic dysfunction and neuron death. Neuron 72, 57–71 (2011).

6 Luk, K. C. et al. Pathological α-synuclein transmission initiates Parkinson-like neurodegeneration in nontransgenic mice. Science 338, 949–953 (2012).

7 Luk, K. C. et al. Intracerebral inoculation of pathological α-synuclein initiates a rapidly progressive neurodegenerative α-synucleinopathy in mice. J Exp. Med. 209, 975–986 (2012).

8 Guo, J. L. & Lee, V. M. Cell-to-cell transmission of pathogenic proteins in neurodegenerative diseases. Nat. Med. 20, 130–138 (2014).

9 Peng, C. et al. Cellular milieu imparts distinct pathological α-synuclein strains in α-synucleinopathies. Nature 557, 558–563 (2018).

10 Riek, R. The Three-Dimensional Structures of Amyloids. Cold Spring Harb. Perspect. Biol. 9 (2017).

11 Fandrich, M., Meinhardt, J. & Grigorieff, N. Structural polymorphism of Alzheimer Aβ and other amyloid fibrils. Prion 3, 89–93 (2009).

12 Li, B. et al. Cryo-EM of full-length α-synuclein reveals fibril polymorphs with a common structural kernel. Nat. Commun. 9, 3609 (2018).

13 Guerrero-Ferreira, R. et al. Two new polymorphic structures of human full-length α-synuclein fibrils solved by cryo-electron microscopy. eLife 8 (2019).

14 Shrivastava, A. N. et al. Differential Membrane Binding and Seeding of Distinct α-Synuclein Fibrillar Polymorphs. Biophys. J (2020).

15 Bousset, L. et al. Structural and functional characterization of two α-synuclein strains. Nat. commun. 4, 2575 (2013).

16 Kim, C. et al. Exposure to bacterial endotoxin generates a distinct strain of α-synuclein fibril. Sci. Rep. 6, 30891 (2016).

17 Ma, M. R., Hu, Z. W., Zhao, Y. F., Chen, Y. X. & Li, Y. M. Phosphorylation induces distinct α-synuclein strain formation. Sci. Rep. 6, 37130 (2016).

18 Lau, A. et al. α-Synuclein strains target distinct brain regions and cell types. Nat. Neurosci. 23, 21–31 (2020).

19 Peelaerts, W. et al. α-Synuclein strains cause distinct synucleinopathies after local and systemic administration. Nature 522, 340–344 (2015).

20 Cremades, N. et al. Direct observation of the interconversion of normal and toxic forms of α-synuclein. Cell 149, 1048–1059 (2012).

21 Paslawski, W., Mysling, S., Thomsen, K., Jorgensen, T. J. & Otzen, D. E. Co-existence of two different α-synuclein oligomers with different core structures determined by hydrogen/deuterium exchange mass spectrometry. Angew. Chem. Int. Ed 53, 7560–7563 (2014).

22 Dusa, A. et al. Characterization of oligomers during α-synuclein aggregation using intrinsic tryptophan fluorescence. Biochemistry 45, 2752–2760 (2006).

23 Ghosh, D. et al. Structure based aggregation studies reveal the presence of helix-rich intermediate during α-Synuclein aggregation. Sci. Rep. 5, 9228 (2015).

24 Serpell, L. C., Fraser, P. E. & Sunde, M. X-ray fiber diffraction of amyloid fibrils. Methods Enzymol. 309, 526–536 (1999).

25 Miake, H., Mizusawa, H., Iwatsubo, T. & Hasegawa, M. Biochemical characterization of the core structure of α-synuclein filaments. J Biol. Chem. 277, 19213–19219 (2002).

26 Kushnirov, V. V., Dergalev, A. A. & Alexandrov, A. I. Proteinase K resistant cores of prions and amyloids. Prion 14, 11–19 (2020).

27 Prusiner, S. B. Prions. Proc. Natl. Acad. Sci. U. S. A. 95, 13363–13383 (1998).

28 Vilar, M. et al. The fold of α-synuclein fibrils. Proc. Natl. Acad. Sci. U. S. A. 105, 8637–8642 (2008).

29 Singh, J. & Udgaonkar, J. B. Unraveling the Molecular Mechanism of pH-Induced Misfolding and Oligomerization of the Prion Protein. J Mol. Biol. 428, 1345–1355 (2016).

30 Moulick, R., Das, R. & Udgaonkar, J. B. Partially Unfolded Forms of the Prion Protein Populated under Misfolding-promoting Conditions: characterization by hydrogen exchange mass spectrometry and nmr. J Biol. Chem. 290, 25227–25240 (2015).

31 Kumar, H., Singh, J., Kumari, P. & Udgaonkar, J. B. Modulation of the extent of structural heterogeneity in α-synuclein fibrils by the small molecule thioflavin T. J Biol. Chem. 292, 16891–16903 (2017).

32 Del Mar, C., Greenbaum, E. A., Mayne, L., Englander, S. W. & Woods, V. L., Jr. Structure and properties of α-synuclein and other amyloids determined at the amino acid level. Proc. Natl. Acad. Sci. U. S. A. 102, 15477–15482 (2005).

33 Schuetz, A. et al. Protocols for the sequential solid-state NMR spectroscopic assignment of a uniformly labeled 25 kDa protein: HET-s(1-227). Chembiochem 11, 1543–1551 (2010).

34 Kumar, R. et al. Cytotoxic Oligomers and Fibrils Trapped in a Gel-like State of α-Synuclein Assemblies. Angew. Chem. Int. Ed 57, 5262–5266 (2018).

35 Gath, J. et al. Unlike twins: an NMR comparison of two α-synuclein polymorphs featuring different toxicity. PLoS One 9, e90659 (2014).

36 Jucker, M. & Walker, L. C. Pathogenic protein seeding in Alzheimer disease and other neurodegenerative disorders. Ann Neurol 70, 532–540 (2011).

37 Buell, A. K. et al. Solution conditions determine the relative importance of nucleation and growth processes in α-synuclein aggregation. Proc. Natl. Acad. Sci. U. S. A. 111, 7671–7676 (2014).

38 Willander, H. et al. BRICHOS domains efficiently delay fibrillation of amyloid ß-peptide. J Biol. Chem. 287, 31608–31617 (2012).

39 Bolognesi, B. et al. ANS binding reveals common features of cytotoxic amyloid species. ACS Chem. Biol. 5, 735–740 (2010).

40 Campioni, S. et al. A causative link between the structure of aberrant protein oligomers and their toxicity. Nat. Chem. Biol. 6, 140–147 (2010).

41 Vermes, I., Haanen, C., Steffens-Nakken, H. & Reutelingsperger, C. A novel assay for apoptosis. Flow cytometric detection of phosphatidylserine expression on early apoptotic cells using fluorescein labelled Annexin V. J. Immunol. Methods 184, 39–51 (1995).

42 Brundin, P., Melki, R. & Kopito, R. Prion-like transmission of protein aggregates in neurodegenerative diseases. Nat. Rev. Mol. Cell Biol. 11, 301–307 (2010).

43 Collinge, J. & Clarke, A. R. A general model of prion strains and their pathogenicity. Science 318, 930–936 (2007).

44 Tu, P. H. et al. Glial cytoplasmic inclusions in white matter oligodendrocytes of multiple system atrophy brains contain insoluble α-synuclein. Ann. Neurol. 44, 415–422 (1998).

45 Volpicelli-Daley, L. A., Luk, K. C. & Lee, V. M. Addition of exogenous α-synuclein preformed fibrils to primary neuronal cultures to seed recruitment of endogenous α-synuclein to Lewy body and Lewy neurite-like aggregates. Nat. protoc. 9, 2135–2146 (2014).

46 Whitt, M. A. & Mire, C. E. Utilization of fluorescently-labeled tetracysteine-tagged proteins to study virus entry by live cell microscopy. Methods 55, 127–136 (2011).

47 Irtegun, S., Ramdzan, Y. M., Mulhern, T. D. & Hatters, D. M. ReAsH/FlAsH labeling and image analysis of tetracysteine sensor proteins in cells. J Vis. Exp. (2011).

48 Ray, S. et al. Liquid-liquid phase separation and liquid-to-solid transition mediate α-synuclein amyloid fibril containing hydrogel formation. bioRxiv, 619858 (2019).

49 Roberti, M. J., Bertoncini, C. W., Klement, R., Jares-Erijman, E. A. & Jovin, T. M. Fluorescence imaging of amyloid formation in living cells by a functional, tetracysteine-tagged α-synuclein. Nat. methods 4, 345–351 (2007).

50 Shen, D. et al. Novel cell-and tissue-based assays for detecting misfolded and aggregated protein accumulation within aggresomes and inclusion bodies. Cell Biochem. Biophys. 60, 173–185 (2011).

51 Guo, J. L. et al. Distinct α-synuclein strains differentially promote tau inclusions in neurons. Cell 154, 103–117 (2013).

52 Uversky, V. N., Li, J. & Fink, A. L. Evidence for a partially folded intermediate in α-synuclein fibril formation. J Biol. Chem. 276, 10737–10744 (2001).

53 Narkiewicz, J., Giachin, G. & Legname, G. In vitro aggregation assays for the characterization of α-synuclein prion-like properties. Prion 8, 19–32 (2014).

54 Cremades, N., Chen, S. W. & Dobson, C. M. Structural Characteristics of α-Synuclein Oligomers. Int. Rev. Cel. Mol. Bio. 329, 79–143 (2017).

55 Petkova, A. T. et al. Self-propagating, molecular-level polymorphism in Alzheimer’s β-amyloid fibrils. Science 307, 262–265 (2005).

56 Shahnawaz, M. et al. Discriminating α-synuclein strains in Parkinson’s disease and multiple system atrophy. Nature 578, 273–277 (2020).

57 Braak, H. et al. Staging of brain pathology related to sporadic Parkinson’s disease. Neurobiol. Aging 24 (2003).

58 Yamasaki, T. R. et al. Parkinson’s disease and multiple system atrophy have distinct α-synuclein seed characteristics. J Biol. Chem. 294, 1045–1058 (2019).

59 Prusiner, S. B. et al. Evidence for α-synuclein prions causing multiple system atrophy in humans with parkinsonism. Proc. Natl. Acad. Sci. U. S. A. 112, E5308–5317 (2015).

60 Volles, M. J. & Lansbury, P. T., Jr. Relationships between the sequence of α-synuclein and its membrane affinity, fibrillization propensity, and yeast toxicity. J Mol Biol. 366, 1510–1522 (2007).

61 Singh, P. K., Kotia, V., Ghosh, D., Mohite, G. M., Kumar, A., and Maji, S. K. Curcumin modulates α-synuclein aggregation and toxicity. ACS Chem. Neurosci. 4, 393–407 (2013).

62 Mehra, S. et al. Glycosaminoglycans have variable effects on α-synuclein aggregation and differentially affect the activities of the resulting amyloid fibrils. J Biol. Chem. 293, 12975–12991 (2018).

63 Jha, N. N. et al. Complexation of NAC-Derived Peptide Ligands with the C-Terminus of α-Synuclein Accelerates Its Aggregation. Biochemistry 57, 791–804 (2018).

64 Singh, J., Sabareesan, A. T., Mathew, M. K. & Udgaonkar, J. B. Development of the structural core and of conformational heterogeneity during the conversion of oligomers of the mouse prion protein to worm-like amyloid fibrils. J Mol. Biol. 423, 217–231 (2012).

65 Singh, J. & Udgaonkar, J. B. Dissection of conformational conversion events during prion amyloid fibril formation using hydrogen exchange and mass spectrometry. J Mol. Biol. 425, 3510–3521 (2013).

66 Gerez, J. A. et al. A cullin-RING ubiquitin ligase targets exogenous α-synuclein and inhibits Lewy body-like pathology. Sci. Transl. Med. 11 (2019).

