## Supplementary information for "α-Synuclein aggregation intermediates form fibril polymorphs with distinct prion-like properties"

**Solid-state NMR spectroscopy (ssNMR) of fibril polymorphs.**

The assignments of Syn fibrils and HMF spectra were performed by the standard backbone walk approach using multiple 3D spectra and relying on the redundancy of chemical shifts. The residues were assigned by starting a walk form threonine or serine residues. These residues have distinct Cα and Cβ chemical shifts and can be easily identified in different spectra. In situation when the walk could not be carried out further the next alanine and threonine peaks were chosen to start a fresh walk. The pairs of residues identified in walk were assigned based on the uniqueness of these pairs in the sequence. The repeating pairs were assigned if only one such pair was unassigned in the sequence. There are few peaks in the Syn fibril that were assigned based on largely similar HMF assignment especially in the 1-40 and 80-98 regions.

Figure S4a shows a plot of the difference in chemical shifts for the different atoms (N, Cα and Cβ) of the α-Syn fibril and HMF samples. The actual assignments are depicted on the 2D $^{13}$C-$^{13}$C and $^{15}$N-$^{13}$C spectra shown in Figure 3a and b of the main text. In both fibril samples, the residues from 1-40 and 81-98 showed negligible chemical shift differences in the N, Cα and Cβ atoms. Across these regions, the average chemical shift difference for the N, Cα and Cβ atoms appeared to be less than 0.2, 0.15 and 0.2 ppm, respectively. Only Cβ atoms for few residues such as 10K 28E, 83E and 92T showed more significant (>0.3 ppm) chemical shift perturbation. This suggests the backbone conformation is identical for the two samples in these regions.

The significant chemical shift differences between the two fibrils forms appeared largely in the segment comprising residues 40-80 (Figure 3a, S4). The peaks in this region generally had weaker intensities but comparable line-widths to other peaks in the spectra. The most likely cause for the weaker peaks could be dynamics in the intermediate regime of structural heterogeneity (61, 62). While it was possible to sequentially assign most residues from 56-80, the lower intensities hampered the sequential backbone assignments (Figure S4a, purple crosses) for the 40-55-residue segment.

After the unambiguous assignment of other regions of the protein, we observed several low-intensity peaks in the spectra for particular residue type pointing to structural heterogeneity. For example, in Figure S4b zoom-in spectra, several Cα-Cβ peaks (marked by black circles) in the alanine region are unassigned; despite only one unassigned alanine residue over the entire sequence (Figure S4a).

For HMF, G41 and G47 peaks were unambiguously assigned while the position of the 44T/54T was guessed assigned based on the fact that all other threonine peaks were already accounted in the backbone assignments (blue spectra, Figure 3a). In the Syn fibril sample, 41/47G and 44/54T could not be assigned while all other threonine and glycine peaks were assigned (red spectra, Figure 3a, zoom-in spectra). We envisage that the weak peaks in the glycine and threonine region belong to the unassigned glycine and threonine residues present in the 40-60 residues segment. From Figure 3a, zoom-in spectra, the differences in peak position for 41/47G and 44/54T residues suggest a difference between the two fibrils forms in the heterogeneous region. Such specific heterogeneous behavior was also observed in the H/D exchange data in main text (Figure 2c). The backbone chemical shift assignment of the HMF and the Syn fibrils closely resemble the assignments with BMRB entry 17498 (1). In this study also, differential scaling of peak intensities was observed in the region 41-60 (1). As shown above, the chemical shift differences in the Cβ atoms were uniformly distributed in the protein and are mainly confined to polar residues (Figure S4b, zoom-in spectra), thereby indicating that it is probably the differences in hydrogen bonding involving the polar side chains or the side-chain packing that lead to observed morphological and biophysical differences in the two fibril polymorphs. The side chain-packing hypothesis is further corroborated by the difference in the diffraction pattern of the two samples in the arrangement of β-sheets along the fibril axis (Figure 1h).

### Figures

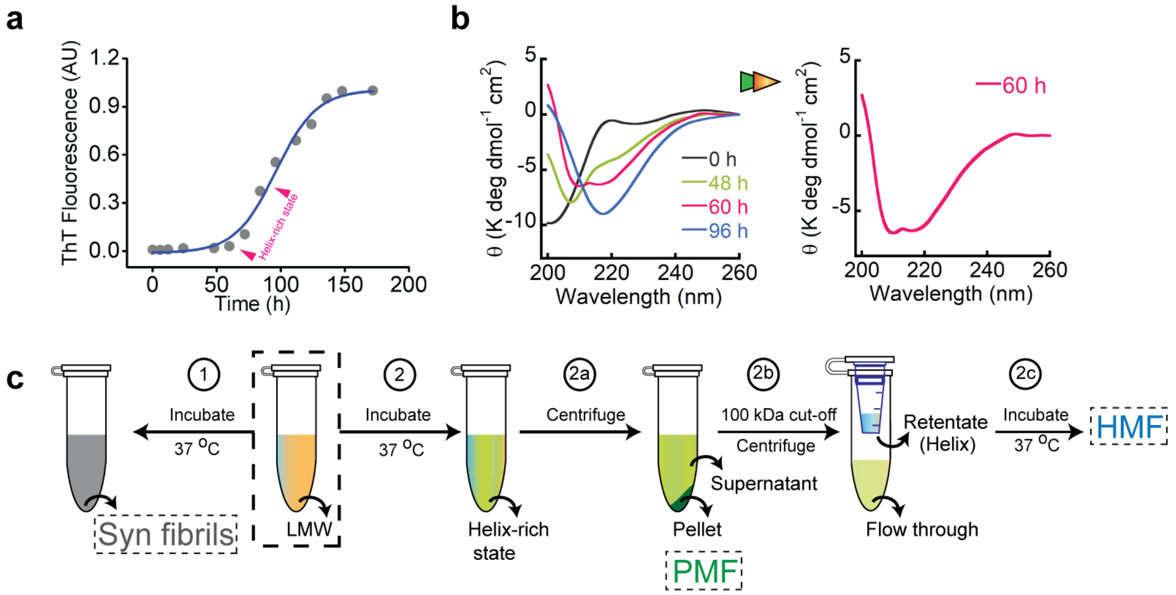

**Figure S1. Isolation of helical-intermediate during α-Syn aggregation.** (a) Kinetics of α-Syn aggregation monitored by amyloid-specific ThT dye and α-helix-rich state (determined by CD) in early elongation phase is marked (pink). (b) Time-dependent structural transition of α-Syn aggregation probed by CD (left) and appearance of helix ~60 h of incubation (right). (c) Schematic for isolating various α-Syn species formed during the aggregation process. Briefly, LMW protein solution is incubated at 37 °C with slight agitation and allowed to mature in fibrils without any isolation of intermediates, termed as Syn fibrils (1). In another set (2), as soon as helix appeared in CD, the solution is centrifuged at 14,000 x g for 30 mins. Pellet fraction is collected and labeled as pre-matured fibrils (PMF). The supernatant after PMF isolation is allowed to pass through 100 kDa cut-off filter and flow through is removed. The retentate (containing helical oligomers) is collected and incubated at 37°C for 8 days for maturation. After incubation, the resulting fibrils are centrifuged to obtain Helix-mature fibrils (HMF).

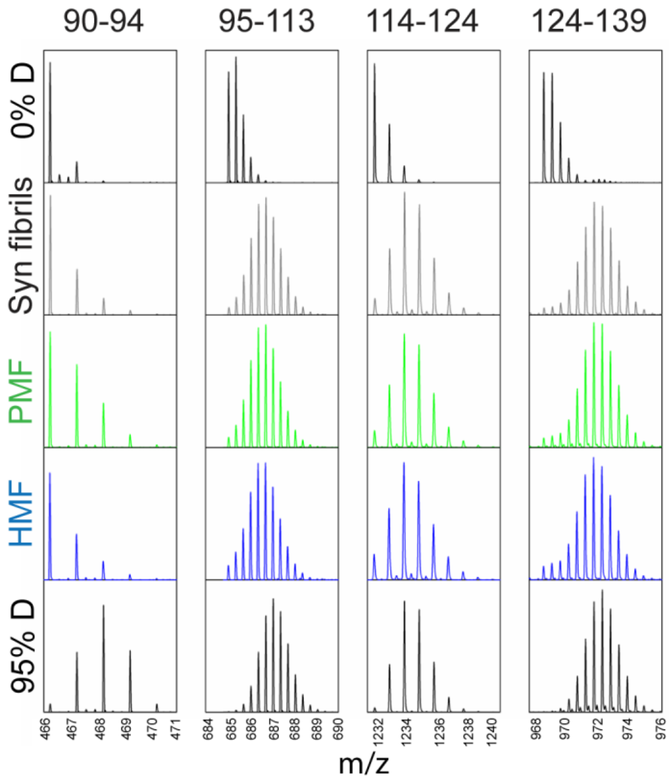

**Figure S2.** Representative mass spectra of peptides of α-Syn from 90-140 sequence (C-terminus region) along with the mass spectra of protonated (0% D) and deuterated (95% D) controls. 1800 sec labeling pulse is given by incubating the fibril samples in deuterated buffer at pH 8.0, 37 $^{o}$C. The sequence spanning residues from 90-140 gets fully labeled in all the fibril polymorphs indicating the flexible nature of C-terminus in the amyloid backbone.

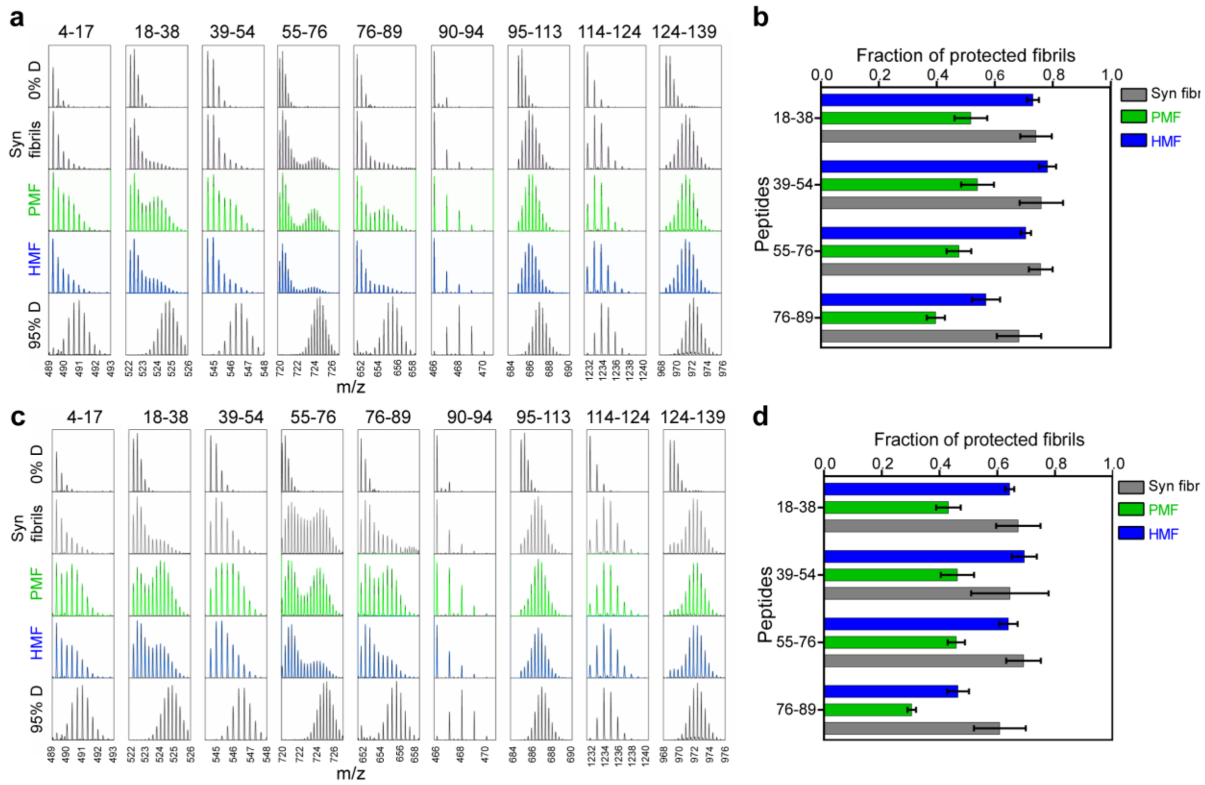

**Figure S3. Hydrogen-Deuterium exchange of fibril polymorphs.** (a) Representative mass spectra of peptide fragments of fibrillar polymorphs at 30 sec labelling pulse in deuterated buffer at pH 7.0, 4 $^{o}$C. The protonated (0% D) and deuterated (95% D) control spectra are shown along with. (b) Bar plot representing the fraction of protected fibril in each polymorph (for residues 18-38, 39-54, 55-76 and 76-89) at 30 sec labelling pulse. Values represent mean ± s.e.m, n=3. (c) Representative mass spectra of peptide fragments of fibrillar polymorphs at 18000 sec labelling pulse in deuterated buffer at pH 8.0, 37 $^{o}$C. The protonated (0% D) and deuterated (95 % D) control spectra are shown along with. (d) Bar plot representing the fraction of protected fibril in each polymorph (for residues 18-38, 39-54, 55-76 and 76-89) at 18000 sec labelling pulse. Values represent mean ± s.e.m, n=3). In all the fibrils polymorphs, sequence spanning residues 4 to 89 remains highly protected and unlabeled, whereas residues 95 to 140 get fully labeled. However, the fraction of protected fibril type in each polymorph is different.

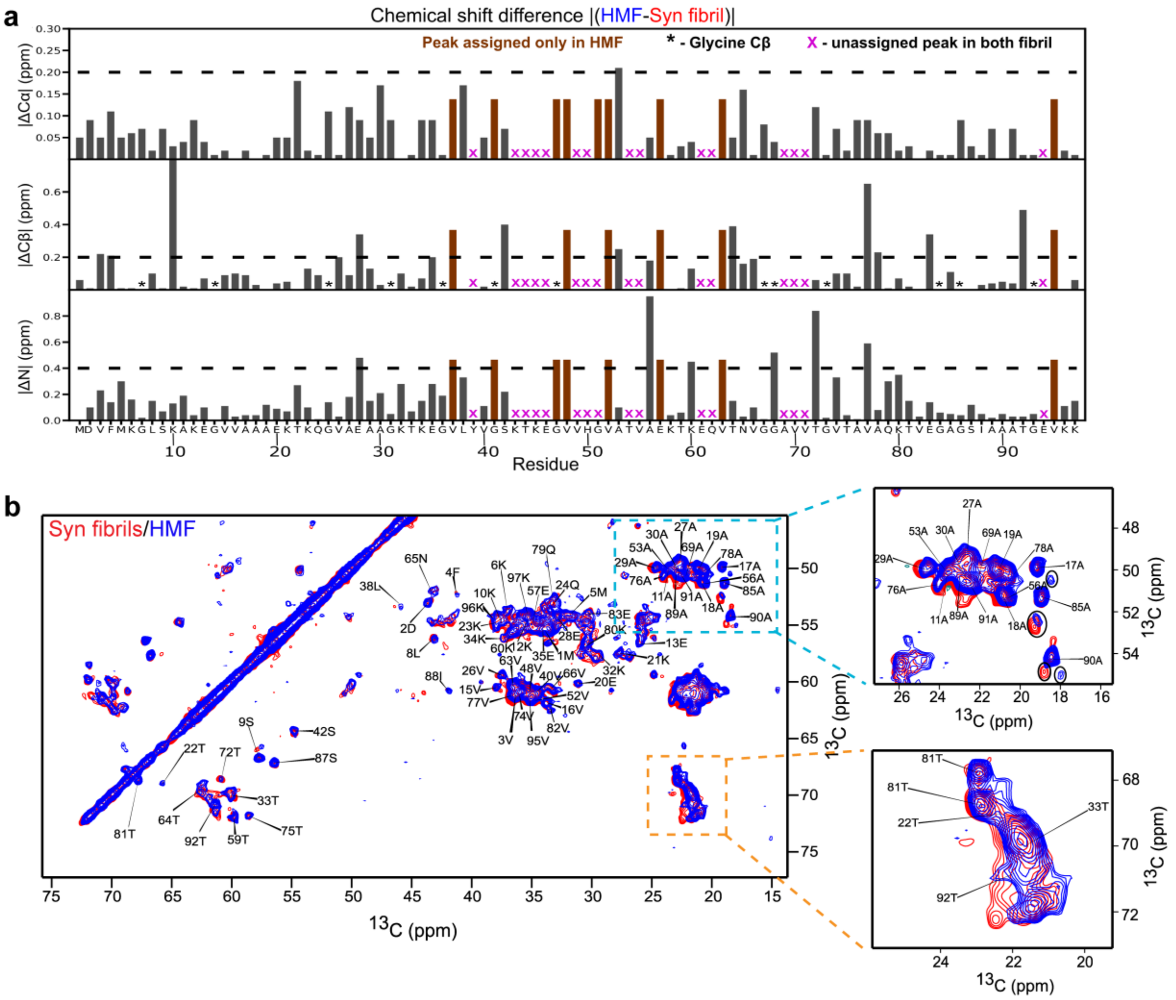

**Figure S4. Solid-state NMR of fibril polymorphs.** Comparison of uniformly $^{13}C,^{15}N$ labeled Syn fibrils (red) with the HMF (blue). (a) The plot shows differences in the values of chemical shift of Syn fibrils and HMF for Cα, Cβ and N. The black dotted line indicates the significant value of change for the respective atom. Brown colour indicates the residues that are unassigned in Syn fibrils. (*) is used to highlight that glycine does not have Cβ atom. Purple X indicates the unassigned residues in both fibril forms. (b) Overlay of 2D DARR of uniformly $^{13}C,^{15}N$ labeled Syn fibrils (red) and HMF (blue). Zoomed in region of the DARR spectrum for alanine and threonine residues are represented alongside. Only the HMF assignments are shown for clarity.

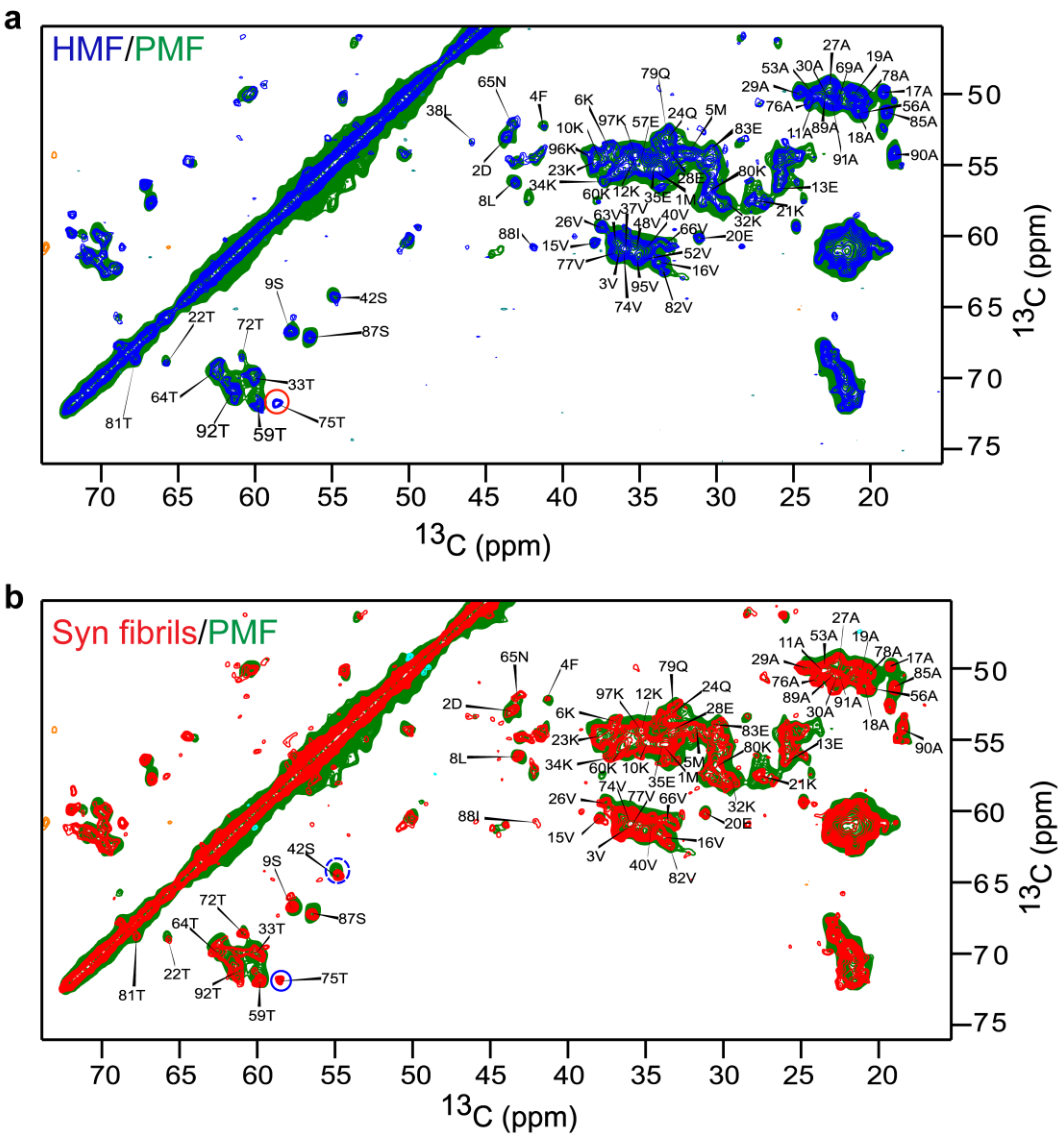

**Figure S5. 2D DARR spectra of fibril polymorphs.** (a) Overlay of 2D DARR of uniformly $^{13}C$,$^{15}N$ labeled HMF (blue) and PMF (green). The assignments of only HMF sample is shown for clarity. (b) Overlay of the 2D DARR spectra of uniformly $^{13}C$,$^{15}N$ labeled Syn fibrils (red) and PMF (green). The assignments depicted are from the Syn fibrils.

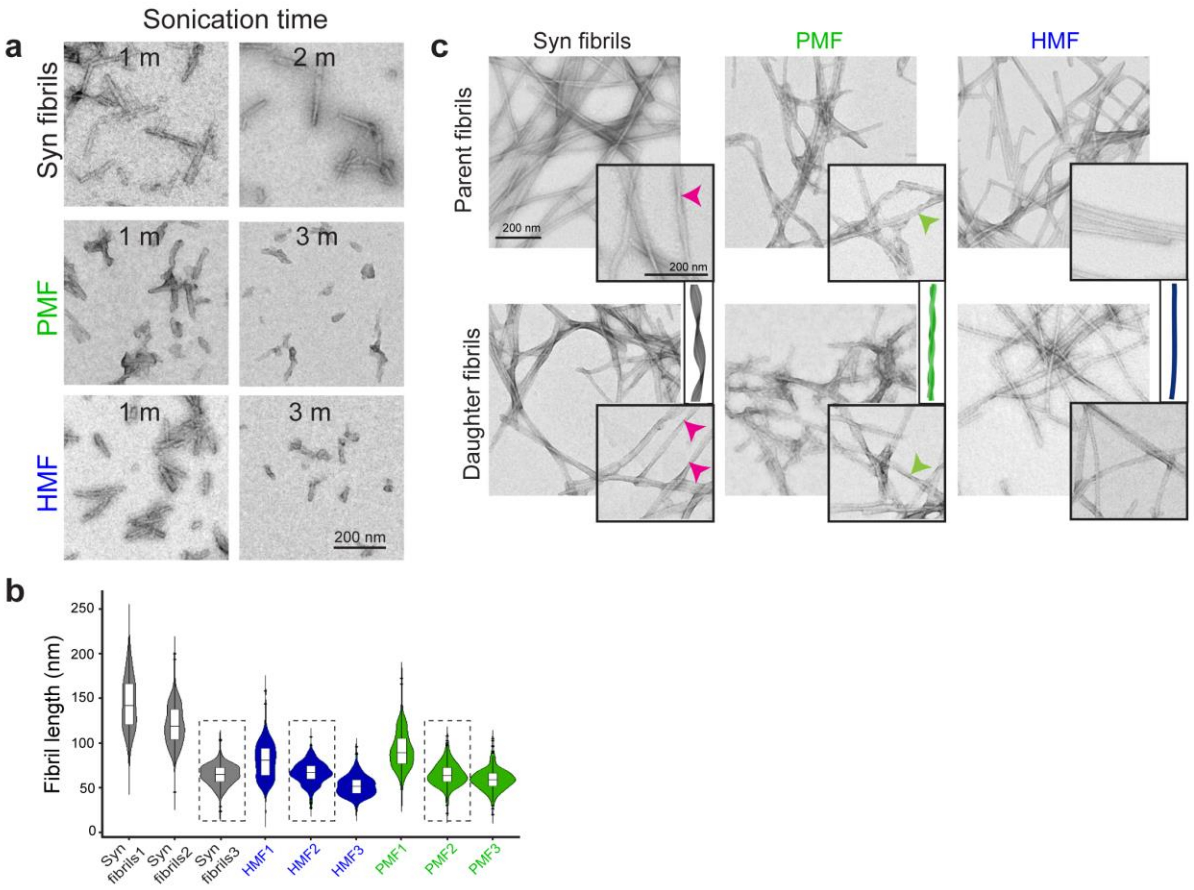

**Figure S6. Seeding ability of fibril polymorphs *in vitro***. (a) Time-dependent optimization of fibrillar seed length. Each fibril sample sonicated for 1 min, 2 mins and 3 mins and TEM images acquired at respective time points are shown. Scale bar represents 200 nm (Syn fibrils; 3 min, HMF and PMF; 2 min is shown in figure 3b of the main text) (b) Violin plot depicts the distribution and the average fibrillar length (nm) post sonication at different time points analyzed by Image J software (n >270 counts). 1, 2 and 3 refer to the sonication time (i.e. 1 min, 2 min and 3 min) in each case. The fibril length of Syn fibrils at 3 min and HMF and PMF obtained at 2 min sonication are nearly same i.e. 65 ± 12 (mean ± s.d.), 65 ± 13 and 66 ± 12 nm, respectively. (c) TEM micrographs of parent fibrils and the fibrils obtained after seeding (daughter fibrils). The inset shows images at higher magnification illustrating the morphological differences amongst the polymorphs that are preserved in the daughter fibrils. Scale bar is 200 nm.

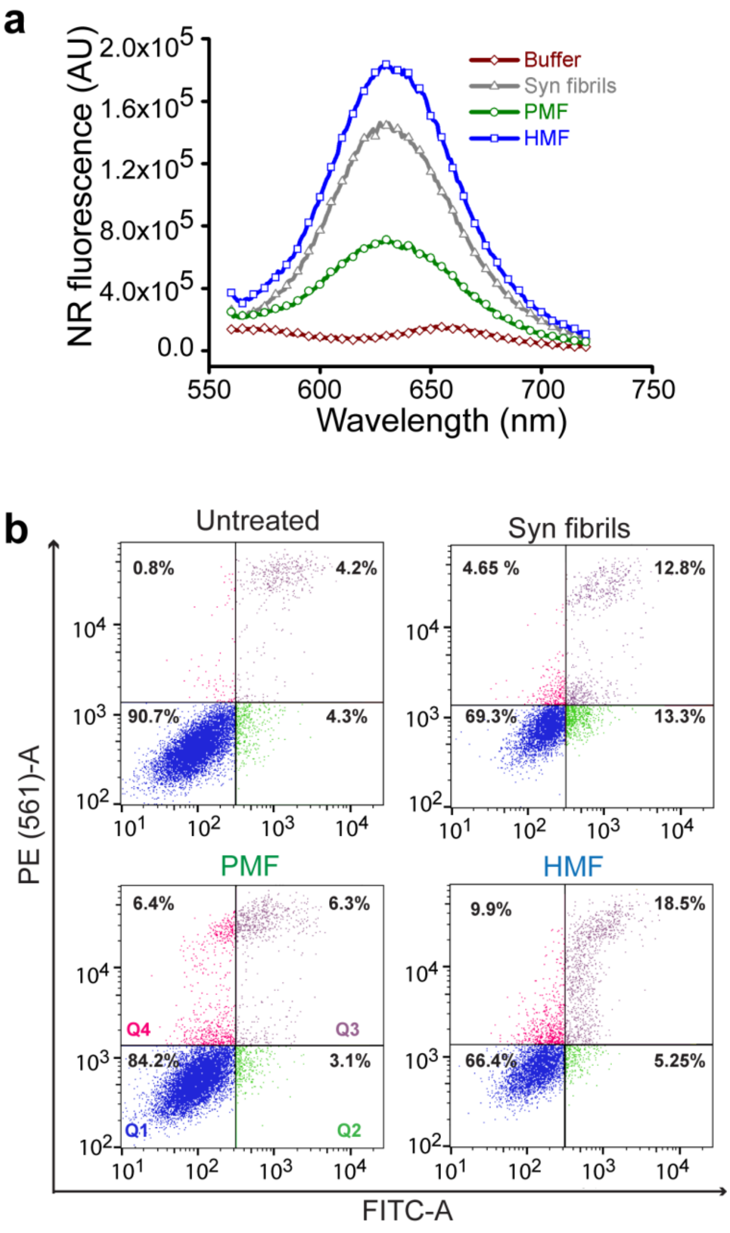

**Figure S7. Hydrophobic exposure and cellular toxicity of fibril polymorphs**. (a) Hydrophobic surface exposure determined by NR binding assay in the wavelength range of 550 to 750 nm. Maximum NR fluorescence is observed in HMF, suggesting the presence of highly exposed hydrophobic surfaces. Cell death analysis of SH-SY5Y cells treated with fibrillar polymorphs (50 µM) for 48 h by Annexin V- PI binding assay using flow cytometry. The colour coding for the four quadrants is Blue (Q1); Viable cells, green (Q2); early apoptotic, purple(Q3); late apoptotic and magenta (Q4); necrotic population. HMF polymorph followed by Syn fibrils shows maximum cell death compared to PMF where ~85% cells are viable. Untreated sample shows the majority (~91 %) of cells as viable and non-apoptotic.

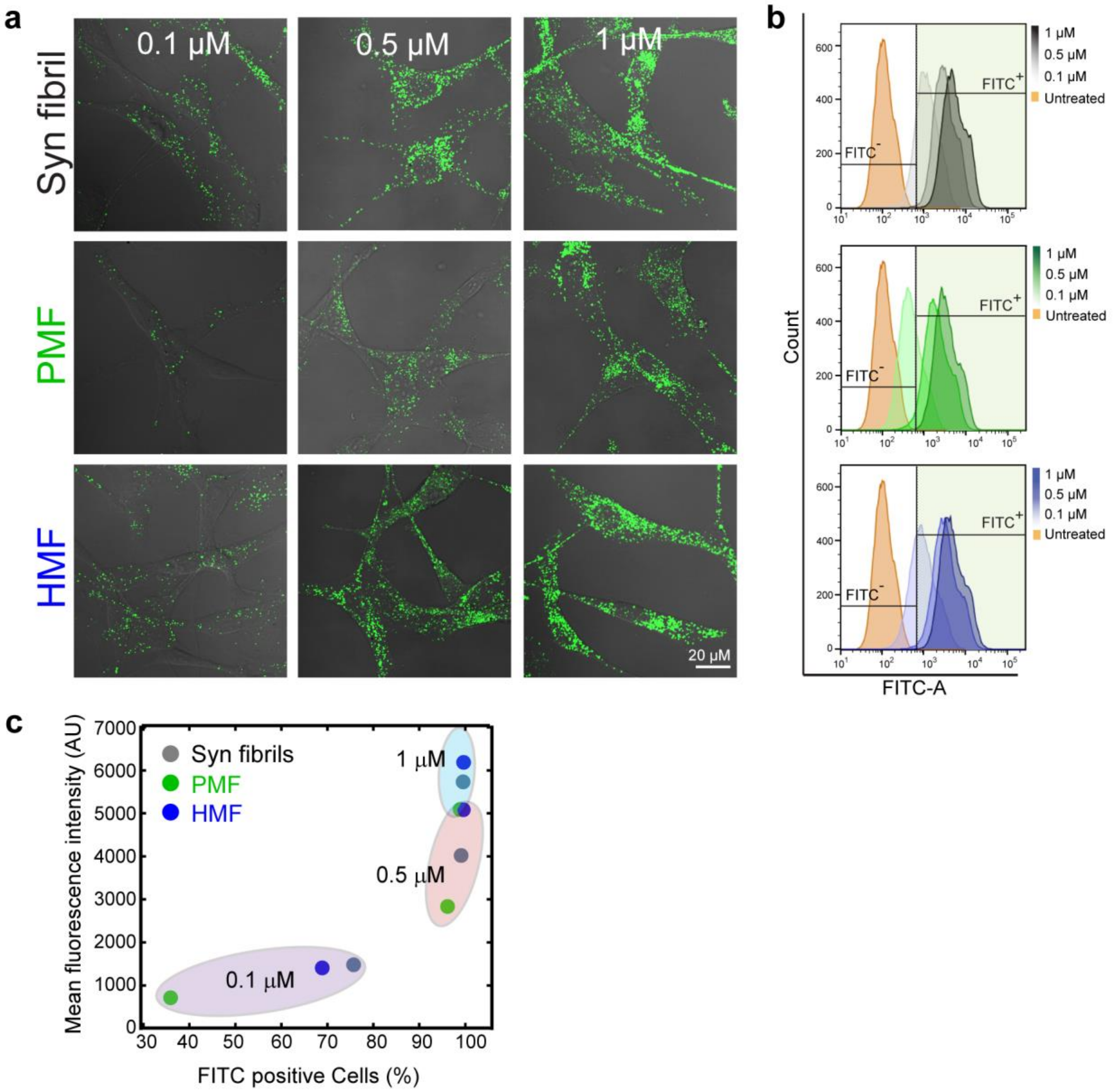

**Figure S8. Internalization ability of α-Syn polymorphs in Glioblastoma cells**. (a) Confocal images of U87-MG cells treated with FITC-labeled fibril polymorphs for 24 h at three different concentrations (0.1 μM, 0.5 μM, 1 μM). Scale bar is 20 μM. (b) Histogram plots depict concentration-dependent increase in the FITC signal in U87-MG cells upon internalization of fibrillar polymorphs along with Syn fibrils. The gating for FITC- (non-shaded area) and FITC+ (shaded area) is based on the fluorescence intensity of untreated sample. (c) The correlation plot of mean fluorescence intensity and percentage of U87-MG cells positive for FITC. The MFI values for each fibril sample are grouped according to their concentration. Cells treated with PMF ≥ 0.5 μM concentration show 80-90 % $FITC^{+}$ cells but the total amount of fibrils internalized (as indicated by respective MFI values) is less than HMF and Syn fibrils.

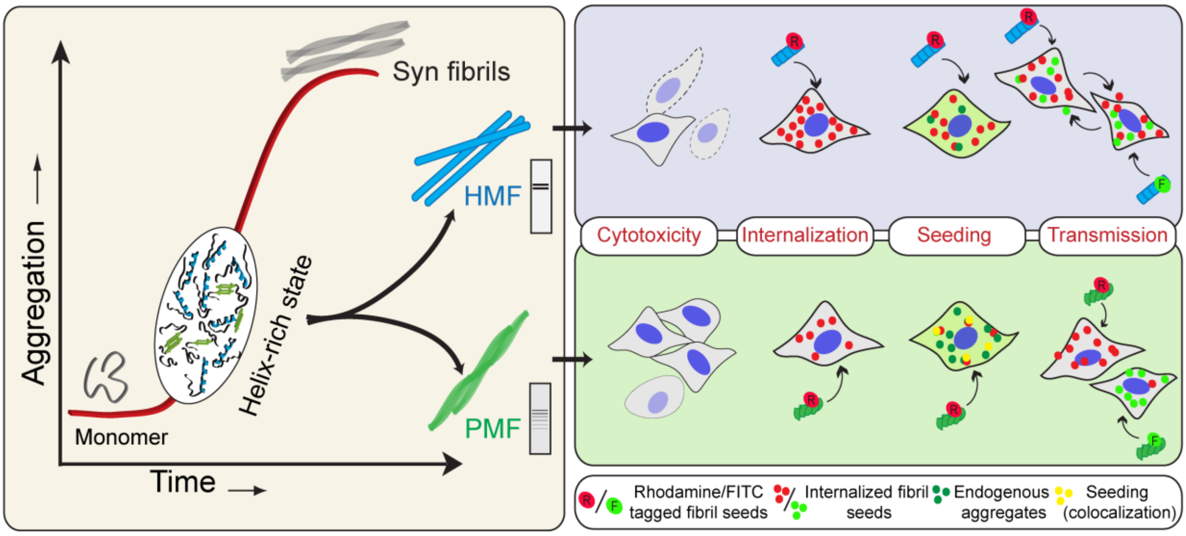

**Figure S9. Aggregation intermediates resulting in fibril polymorphs with distinct prion-like behavior** (a) Schematic representation of α-Syn fibril polymorphs generated from the helix-rich state during aggregation and comparative analysis of their functional properties. HMF is potentially toxic polymorph and is readily uptaken by cells and exhibits cell-to-cell transmission ability. On the contrary, PMF lacks these properties but exhibits high seeding potency in cells.

**Supplementary Table 1**. The experimental parameters used for ssNMR.

| **Sample** | **Syn Fibril** | | **PMF** | | **HMF** | |
|---|---|---|---|---|---|---|
| Experiment | DARR | NCA | DARR | NCA | DARR | NCA |
| Spectrometer 1H frequency | 700 MHz | 700 MHz | 700 MHz | 700 MHz | 700 MHz | 700 MHz |
| Probe | 4mm Bruker | 4mm Bruker | 4mm Bruker | 4mm Bruker | 4mm Bruker | 4mm Bruker |
| MAS (kHz) | 12.5 | 12.5 | 12.5 | 12.5 | 12.5 | 12.5 |
| Measurement time | 26h | 6h | 46h | 20h | 19h | 7h |
| Number of scans | 28 | 128 | 90 | 400 | 20 | 160 |
| Pulse delays (s) | 2.5 | 2.5 | 2.5 | 2.5 | 2.5 | 2.5 |
| Transfer 1 | HC-CP | HN-CP | HC-CP | HN-CP | HC-CP | HN-CP |
| Time (ms) | 0.6 | 0.9 | 0.7 | 1 | 0.5 | 1.5 |
| RF field (kHz) | 60.86(H)/43.8(C) | 55.23(H)/38.4(N) | 66.23(H)/44.6(C) | 50.29(H)/40.3(N) | 62.93(H)/43.8(C) | 56.72(H)/39(N) |
| Shape | tangent | tangent | tangent | tangent | tangent | tangent |
| Carrier (ppm) | 89.2 | 112 | 89.2 | 112 | 89.2 | 113 |
| Transfer 2 | DARR | NCA-CP | DARR | NCA-CP | DARR | NCA-CP |
| Time (ms) | 35 | 5 | 35 | 5 | 35 | 5 |
| RF field (kHz) | 12.5(H) | 7.37(C)/4.84(N) | 12.5(H) | 7.72(C)/5.08(N) | 12.5(H) | 7.71(C)/5.03(N) |
| Shape | - | tangent | - | tangent | - | tangent |
| Carrier (ppm) | - | 49.6 | - | 49.6 | - | 49.6 |
| t1 increments | 1202 | 64 | 710 | 72 | 1250 | 64 |
| SW (ppm) | 250 | 35 | 250 | 40 | 250 | 32 |

| | | | | | | |
|---|---|---|---|---|---|---|
| Carrier (ppm) | 89.2 | 112 | 89.2 | 112 | 89.2 | 113 |
| Acquisition time (ms) | 13.7 | 12.8 | 8.1 | 12.6 | 14.2 | 14 |
| t2 increments | 1778 | 1696 | 1536 | 1776 | 1776 | 1778 |
| SW (ppm) | 338 | 338 | 338 | 338 | 338 | 3338 |
| Carrier (ppm) | 89.2 | 49.6 | 89.2 | 49.6 | 89.2 | 49.6 |
| Acquisition time (ms) | 14.9 | 14.2 | 12.9 | 14.9 | 14.9 | 14.9 |
| Decoupling | $rCW^{ApA}$ | $rCW^{ApA}$ | $rCW^{ApA}$ | $rCW^{ApA}$ | $rCW^{ApA}$ | $rCW^{ApA}$ |
| RF field (kHz) | 74.4 | 74.4 | 86.2 | 86.2 | 79.3 | 79.3 |

**Supplementry Table 2**. The list of primers used for qRT-PCR.

| Genes | Primers [Forward (F) 5’- 3’ and Reverse (R) 3’- 5’] |
|---|---|
| GAPDH | F: CATTTTACGCTGATCCAGG |
| | R: GGGTTCGAAATGAGGATG |
| p53 | F: ACCTATGGAAACTACTTCCTGAAAA |
| | R: CCGGGGACAGCATCAAATCA |
| Bax | F: AACTGGACAGTAACATGGAG |
| | R: TTGCTGGCAAAGTAGAAAAG |
| p21 | F: AAGACCATGTGGACCTGT |
| | R: GGTAGAAATCTGTCATGCTG |
| Bcl-2 | F: CTGCACCTGACGCCCTTCACC |
| | R: CACATGACCCCACCGAACTCAAAGA |
| Bcl-XL | F: GATCCCCATGGCAGCAGTAAAGCAAG |
| | R: CCCCATCCCGGAAGAGTTCATTCACT |
| Caspase-3 | F: ACATGGAAGCGAATCAATGGACTC |

| | |
|---|---|
| | R: AAGGACTCAAATTCTGTTGCCACC |
| Caspase-9 | F: GCTCTTCCTTTGTTCATC |

References

1. Gath J, Habenstein B, Bousset L, Melki R, Meier BH, Bockmann A. Solid-state NMR sequential assignments of alpha-synuclein. Biomolecular NMR assignments. 2012;6(1):51-5.
